# Molecular impact of nicotine and smoking exposure on the developing and adult mouse brain

**DOI:** 10.1101/2024.11.05.622149

**Authors:** Daianna Gonzalez-Padilla, Nicholas J. Eagles, Marisol Cano, Geo Pertea, Andrew E. Jaffe, Kristen R. Maynard, Dana B. Hancock, James T. Handa, Keri Martinowich, Leonardo Collado-Torres

## Abstract

Maternal smoking during pregnancy (MSDP) is associated with significant cognitive and behavioral effects on offspring. While neurodevelopmental outcomes have been studied for prenatal exposure to nicotine, the main psychoactive component of cigarette smoke, its contribution to MSDP effects has never been explored. Comparing the effects of these substances on molecular signaling in the prenatal and adult brain may provide insights into nicotinic and broader tobacco consequences that are developmental-stage specific or age-independent. Pregnant mice were administered nicotine or exposed to chronic cigarette smoke, and RNA-sequencing was performed on frontal cortices of postnatal day 0 pups born to these mice, as well as on frontal cortices and blood of the adult dams. We identified 1,010 and 4,165 differentially expressed genes (DEGs) in nicotine and smoking-exposed pup brains, respectively (FDR<0.05, Ns = 19 nicotine-exposed vs 23 vehicle-exposed; 46 smoking-exposed vs 49 controls). Prenatal nicotine exposure (PNE) alone was related to dopaminergic synapses and long-term synaptic depression, whereas MSDP was associated with the SNARE complex and vesicle transport. Both substances affected SMN-Sm protein complexes and postsynaptic endosomes. Analyses at the transcript, exon, and exon-exon junction levels supported gene level results and revealed additional smoking-affected processes. No DEGs at FDR<0.05 were found in adult mouse brain for any substance (12 nicotine-administered vs 11 vehicle-administered; 12 smoking-exposed vs 12 controls), nor in adult blood (12 smoking-exposed vs 12 controls), and only 3% and 6.41% of the DEGs in smoking-exposed pup brain replicated in smoking-exposed blood and human prenatal brain, respectively. Together, these results demonstrate variable but overlapping molecular effects of PNE and MSDP on the developing brain, and attenuated effects of both smoking and nicotine on adult versus fetal brain.

## INTRODUCTION

As of 2021, 4.6% of mothers in the United States smoked cigarettes during pregnancy. Although declining in prevalence over time, maternal smoking during pregnancy (MSDP) remains a major public health problem due to the risk it imposes on the health of hundreds of thousands of mothers and their offspring (1,2). Adverse health implications for pregnant women include increased risk for preterm deliveries and miscarriages, and impacts on lung and brain development from various toxic compounds in tobacco smoke for the unborn (3). Prenatal tobacco exposure is also associated with cognitive and behavioral disruption. Specifically, exposed babies are predisposed to impaired language and learning skills, attention deficits, conduct and behavioral alterations, and are at higher risk of developing substance use disorders (4). Several studies have investigated MSDP and prenatal nicotine exposure (PNE) in animal models and confirmed similar effects (4).

Because cigarette smoke contains a mixture of over 7,000 compounds (5), understanding the molecular mechanisms and cellular processes by which tobacco smoke affects neurodevelopment is complex. Many of these constituents are toxic or carcinogenic, and can disrupt brain function (6–10). However, little information is available regarding how individual components of cigarette smoke affect the developing brain during prenatal exposure. The most comprehensively studied substance is nicotine, the main psychoactive component of cigarette smoke. Nicotine activates and desensitizes nicotinic acetylcholine receptors (AChRs) in the developing central nervous system (CNS), and impacts brain development (4,11). Despite extensive data demonstrating a causal association between PNE and brain function (11), the extent to which PNE accounts for the effects of MSDP is not known. However, identifying the molecules and pathways driven by nicotine versus other components present in tobacco smoke is critical to understand impacts of MSDP on neurodevelopment. A transcriptomic investigation of the human prefrontal cortex from postmortem brain donors identified 14 MSDP-associated differentially expressed genes, but did not specifically assess effects of nicotine versus other substances (12). Model organisms can be useful to further study MSDP in controlled settings to untangle nicotine-specific contributions.

Here, we investigated molecular impacts of prenatal exposure in mice of both chronic cigarette smoke and nicotine on offspring (P0: postnatal day 0) as well as to the adult, exposed females compared to controls. Differential expression analysis of frontal cortex tissue revealed changes at the gene level when comparing exposed and unexposed pup brain samples. Affected features by prenatal nicotine and smoking exposure were contrasted and were compared against changes observed in adult brain. These results overlap with previous reports in human, identifying several convergent gene targets. Together, the findings suggest differential, but overlapping transcriptomic modifications from gestational exposure to nicotine and cigarette smoke on the developing brain. Novel PNE and MSDP-associated changes were identified in expression features beyond gene expression modifications, and variability in differential gene expression due to tobacco exposure (nicotine and cigarette smoke) across age (prenatal or adult brain), tissue (brain or blood), and species (human or mouse brain) was noted.

## MATERIALS AND METHODS

Detailed materials and methods can be found in **Supplementary Materials and Methods**.

### Samples

The frontal cortex was isolated from P0 offspring and adult females that delivered the pups, across two separate experiments: one for gestational smoking, and one for nicotine administered during gestation. Blood samples were collected from all smoking-exposed and control adults. In total, 208 samples were collected: 184 brain samples and 24 blood samples (**Fig. 1A, Table S1**). We isolated total RNA and performed bulk RNA sequencing (**Fig. 1B**; **Supplementary Materials and Methods**).

**Figure 1:**
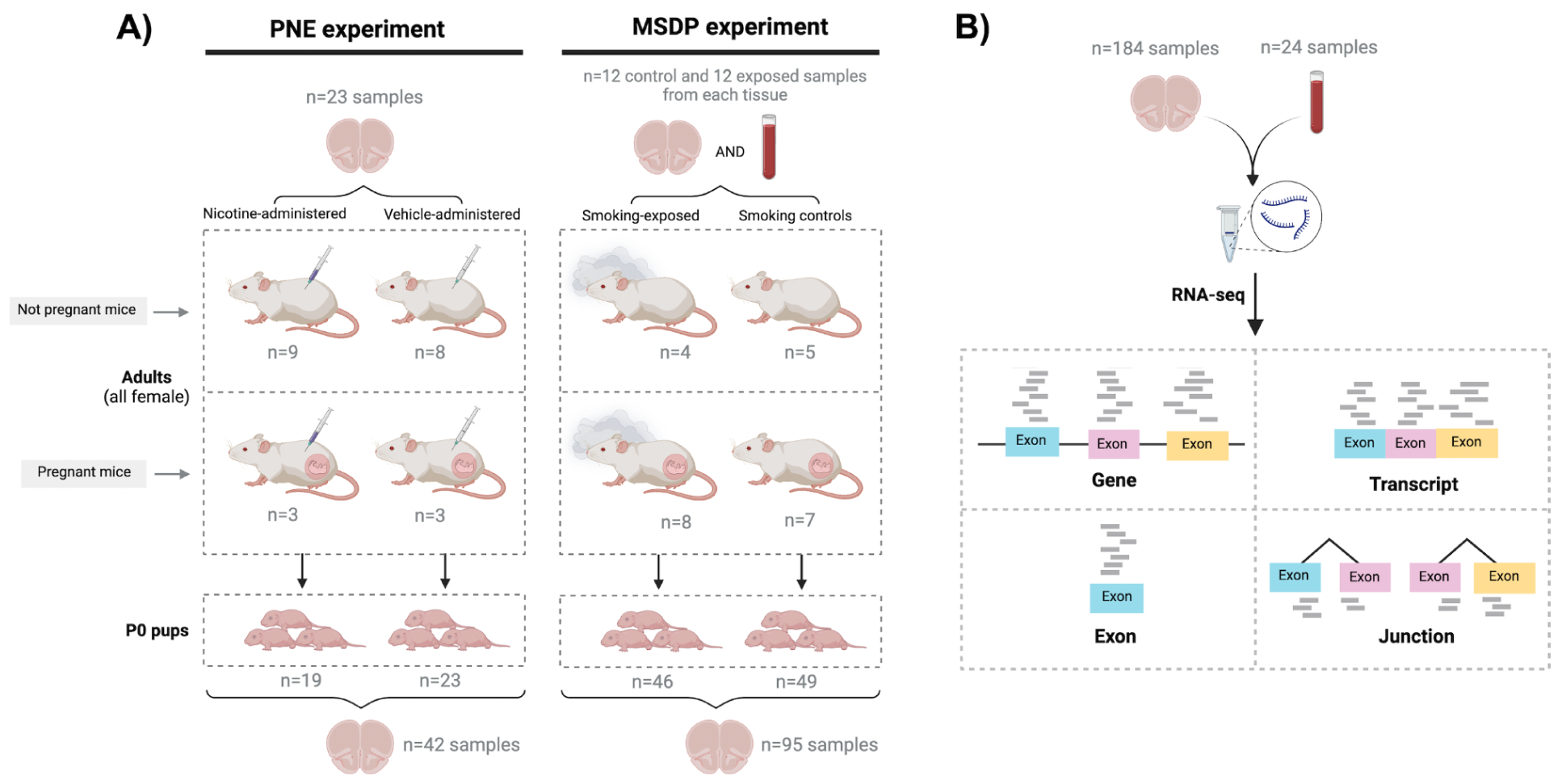
Experimental design of the study. **A)** 21 pregnant mice were split into two experiments: in the first one prenatal nicotine exposure (PNE) was modeled administering nicotine (n=3) or vehicle (n=3) to the dams during gestation, and in the second maternal smoking during pregnancy (MSDP) was modeled exposing dams to cigarette smoke during gestation (n=8) or using them as controls (n=7). A total of 137 pups were born: 19 were born to nicotine-administered mice, 23 to vehicle-administered mice, 46 to smoking-exposed mice, and 49 to smoking control mice. 17 nonpregnant adult females were also nicotine-administered (n=9) or vehicle-administered (n=8) to model adult nicotine exposure, and 9 additional nonpregnant dams were smoking-exposed (n=4) or controls (n=5) to model adult smoking. Frontal cortex samples of all P0 pups (n=137: 42 for PNE and 95 for MSDP) and adults (n=47: 23 for the nicotine experiment and 24 for the smoking experiment) were obtained, as well as blood samples from the smoking-exposed and smoking control adults (n=24), totaling 208 samples. Number of donors and samples are indicated in the figure. **B)** RNA was extracted from such samples and bulk RNA-seq experiments were performed, obtaining expression counts for genes, transcripts, exons, and exon-exon junctions.

### RNA-seq data processing and exploration

Raw sequencing reads were pre-processed and aligned with *SPEAQeasy* (13) and used for expression quantification of genes, transcripts, exons, and exon-exon junctions (**Fig. 1B**). After normalizing read counts and filtering out lowly-expressed features (**Fig. S1**), samples were separated by tissue and age (**Fig. S2**, **Fig. S3**) and filtered by quality control metrics (**Fig. S4**). Dimensionality reduction analysis identified poor-quality samples that were further removed (**Fig. S5**, **Fig. S6**), and revealed transcriptomic sample differences driven by experiment among adult brains (**Fig. S5**, **Fig. S7A**, **Fig. S8**), by sex among pup brains (**Fig. S6**, **Fig. S7B, Fig. S9**), and by pregnancy in blood (**Fig. S3B, Fig. S10**). After discarding poor-quality samples, 23 blood samples, 39 adult brain samples, and 130 pup brain samples were used (**Table S2**).

Additional sample-level sources of gene expression variation were identified through variance partition and canonical correlation analyses, which informed the design of the statistical models used for differential expression analysis (DEA) (**Fig. S11**, **Fig. S12**).

### Differential Expression Analysis (DEA)

Five differential gene expression analyses were performed under the empirical Bayesian framework of *limma*-*voom* (14), comparing 1) nicotine vs vehicle exposure in pup brain, 2) smoking exposure vs control in pup brain, 2) nicotine vs vehicle administration in adult brain, 4) smoking exposure vs control in adult brain, and 5) smoking exposure vs control in adult blood (**Fig. S13**). Gene expression was adjusted for quality control metrics and batch effects, and by sex in pup brain, and pregnancy in adult brain and blood. DEA of expression features other than genes were performed for smoking and nicotine exposures in pup brain (**Fig. S1**). Only genes, transcripts, and exon-exon junctions with *p*-values adjusted for a false discovery rate (FDR) <5%, as well as exons with an FDR<5% and |log2FC|>0.25, were considered differentially expressed (DE).

Resulting moderated gene *t*-statistics were compared between experiments, ages, tissues, and against results from a previous transcriptomic study of prenatal and adult smoking exposure in human dorsolateral prefrontal cortex (12) (**Fig. S1**).

### Functional enrichment analysis

Genes annotated in Gene Ontology (GO) terms and in pathways of the Kyoto Encyclopedia of Genes and Genomes (KEGG), were assessed for their enrichment among our sets of genes applying one-sided Fisher’s exact tests, as implemented in *clusterProfiler* (15), and were FDR controlled.

## RESULTS

The frontal cortex was isolated from P0 offspring across two separate experiments: 1) pups born to female mice exposed to gestational smoking (n=46) or pups born to control female mice (n=49); 2) pups born to female mice administered nicotine during gestation (n=19) or pups born to female mice administered vehicle during gestation (n=23). The frontal cortex was also collected from the adult females that delivered the pups plus additional nonpregnant dams that were: 1) exposed to cigarette smoke (n=12; 8 pregnant) or smoking controls (n=12; 7 pregnant), and 2) administered nicotine (n=12; 3 pregnant) or vehicle-administered (n=11; 3 pregnant). Additionally, blood samples were collected from all smoking-exposed and control adults (n=24, **Fig. 1A**, **Table S1**). We isolated total RNA from all 208 samples and performed bulk RNA sequencing. From these data, we measured the transcriptome at four expression feature levels: genes, transcripts, exons, and exon-exon junctions (**Fig. 1B**). Poor-quality samples were discarded, resulting in a final study size of 130 pup brain samples, 39 adult brain samples, and 23 blood samples (n=192, **Materials and Methods**, **Table S2**, **Fig. S1**).

### Molecular impact of gestational exposure to nicotine and smoking on developing frontal cortex of offspring

From the frontal cortex of P0 offspring from both the prenatal nicotine exposure (PNE) and maternal smoking during pregnancy (MSDP) experiments, we performed differential expression analysis (DEA) at gene, transcript, exon, and exon-exon junction levels (**Fig. 1**, **Fig. S1**). Expression features other than genes were analyzed to support and complement gene-level inferences (16–18). Biologically, gene-level expression is composed by adding transcript-level expression, although gene-level RNA-seq quantification is performed by different computational methods (16,19,20). The highest expressed transcript in a gene can dominate gene-level expression measurements, masking out transcript-level changes (16–18). In addition, transcripts of the same gene with opposing expression directionalities can cancel each other out (16–18). Moreover, exons and exon-exon junction counts can provide additional insights into transcript abundances and alternative splicing (21–24).

Comparing nicotine to vehicle exposure (PNE experiment), we identified 1,010 differentially expressed genes (DEGs, FDR<0.05) (**Fig. 2A**, **Table S3**); 280 DEGs were downregulated and 730 were upregulated. The top two most significantly up- and down-regulated genes were *Foxn3* and *Arrdc3,* and *Senp8* and *Coa4*, respectively (**Fig. 2A**). Comparing smoking exposure to control (MSDP experiment), 4,165 genes were differentially expressed (FDR<0.05): 2,106 were downregulated and 2,059 upregulated (**Fig. 2B**, **Table S4**). *Top2a* and *Tpx2* were the most significant DEGs and were downregulated, followed by the upregulated *AB041806* and *Mt2* genes (**Fig. 2B**). While differential gene expression (DGE) results were poorly correlated between the two experiments (rho=0.13, **Fig. 2C**), we identified 187 shared upregulated genes and 35 shared downregulated genes (**Fig. 2C**). Among the shared upregulated DEGs, *Strap, Snrpd3*, and *Snrpb* act in SMN-Sm protein complexes and *Nsg1*, *Clstn1*, and *Rab4a* in postsynaptic endosomes (**Fig. S14A**, **Fig. S15A,B**). Additionally, 496 genes were upregulated after nicotine exposure, but were not affected by cigarette smoke (**Fig. 2C**), of which 15 were associated with dopaminergic synapses and 8 with long-term synaptic depression (**Fig. S14B**, **Fig. S15C,D**). Similarly, 1,855 genes were upregulated after smoking exposure, but unaltered by nicotine (**Fig. 2C**), with 15 genes involved in the SNARE complex (**Fig. S14A**, **Fig. S15E**), which mediates neurotransmitter release. Furthermore, 17 DEGs were upregulated by smoking exposure and downregulated by nicotine exposure (**Fig. 2C**); of these, *Stx17* and *Bnip1* were enriched for the SNARE complex (**Fig. S14A-C**, **Fig. S15F**). 47 genes were upregulated by nicotine and downregulated by smoking (**Fig. 2C**), 4 showing enrichment for heat shock protein binding activity (**Fig. S14C**, **Fig. S15G**). A summary of the DGE results is provided in **Table S5**.

**Figure 2:**
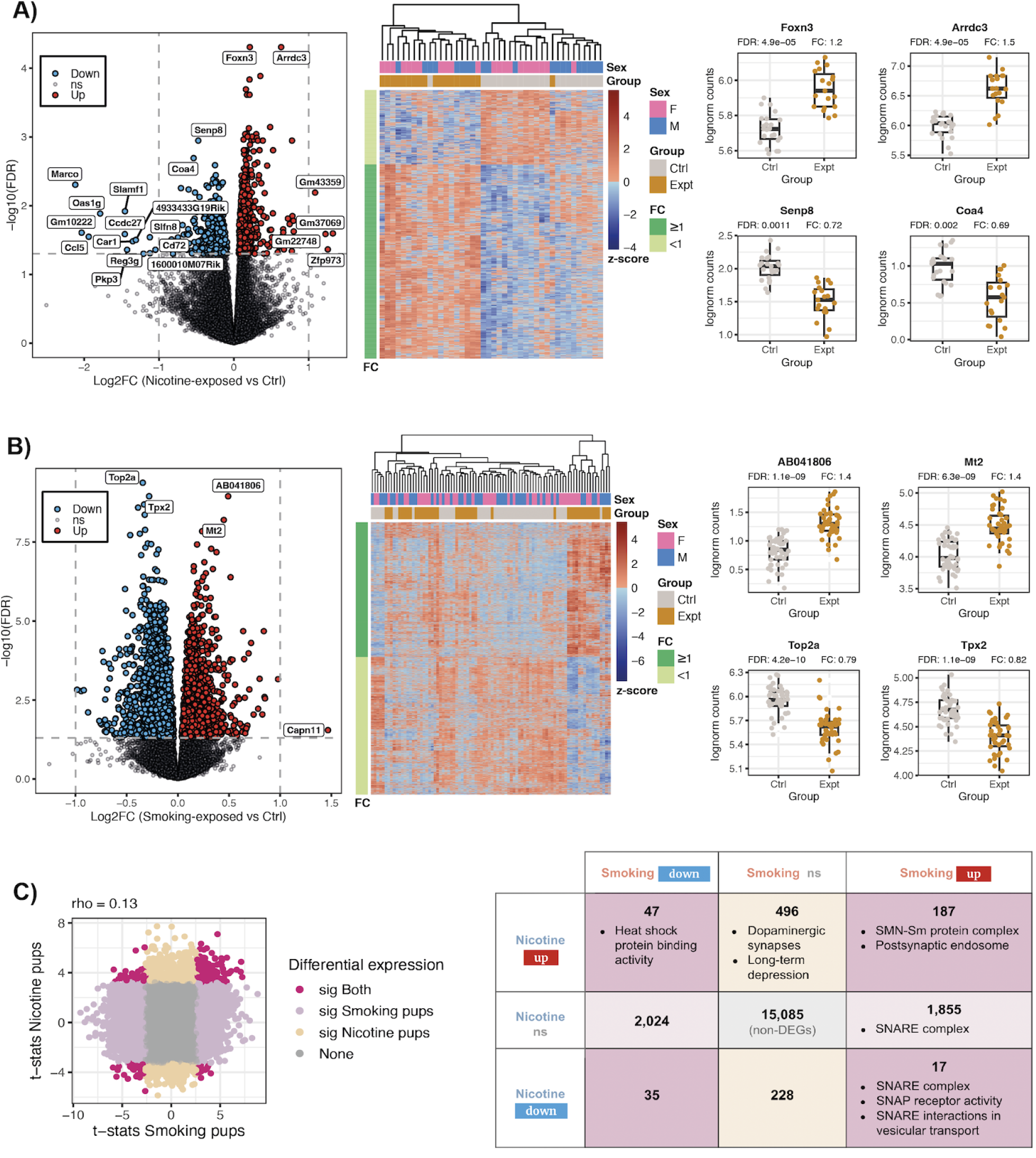
Differentially expressed genes in pup brain. Results of the differential gene expression analysis for **A)** prenatal nicotine vs vehicle exposure (PNE experiment) and **B)** prenatal smoking exposure vs control (MSDP experiment): volcano plots (left) show for each gene its log2-fold-change (logFC) and the -log10 of its false discovery rate (FDR) adjusted *p*-value for differential expression; in blue the DEGs (FDR<0.05) that were downregulated and in red the ones that were upregulated; non-significant (ns) genes appear in gray; labeled genes had |logFC|>1 or were the top 2 most significantly up- or down-regulated genes. Heat maps (middle) show the z-scores for the log2-CPM of the DEGs across samples; left color bars show the FC direction of the genes and top color bars the corresponding sex and experimental group of the samples. Box plots (right) show the log2-CPM of the top 2 most significant up- and down-regulated DEGs in control (Ctrl) and exposed samples (Expt). **C)** Scatter plot of the moderated *t*-statistics for differential expression of the genes for smoking and nicotine exposure. In dark pink the genes that were significantly DE under both exposures, in light pink and beige the ones that were significant for smoking or nicotine exposure only, respectively, and in gray genes that were not significant in any of the experiments; rho corresponds to the Spearman correlation coefficient. The right table presents the number of up- and down-regulated DEGs, as well as non-significant genes, for both nicotine and smoking exposures in pup brain. The molecular functions, cellular components and pathways that are significantly enriched in the given sets of DEGs are indicated. Related to **Fig. S14**, **Fig. S15**, **Table S3**, **Table S4** and **Table S5**.

Differential transcript expression (DTE) analysis identified 232 DE transcripts (FDR<0.05, mapping to 220 unique genes) for nicotine versus vehicle exposure (**Table S6**) and 4,059 DE transcripts (mapping to 3,451 unique genes) for smoking exposure versus control (**Table S7**, **Fig. S16**). Comparing DTE against DGE results for nicotine exposure, DE statistics were concordant at the gene and transcript levels (rho=0.41, **Fig. 3A**, **Table S8**), and most transcripts of DEGs were not differentially expressed, reflecting the transcript diversity for each gene. Similarly, for smoking exposure gene- and transcript-level DE statistics were concordant (rho=0.50, **Fig. 3A**, **Table S9**). However, some genes such as *Phf3, Ankrd11*, *Trpc4*, *Bcl11a*, *Scaf11, Dgcr8*, *Pnsir,* and *Dcun1d5* for nicotine exposure, and *Btf3*, *Cyhr1*, *H13*, *Srsf6, Meaf6*, *Ivns1abp*, *Morf4l2*, *Sin3b*, and *Ppp2r5c* for smoking exposure presented dissimilar DTE and DGE results (**Fig. S17**). Contrasting DTE results across exposures (**Fig. S16**), functional gene profiles for the DE transcripts corroborated and expanded DGE results (**Fig. S14**). We found 1,427 genes expressing upregulated transcripts under smoking exposure only, including 14 and 55 genes encoding for proteins associated with the SNARE complex and transport vesicles, respectively (**Fig. S14E**; **Fig. S15E,H**), as well as 68 genes involved in Parkinson’s, Huntington’s, or prion-related diseases (**Fig. S14F**, **Fig. S15I**).

**Figure 3:**
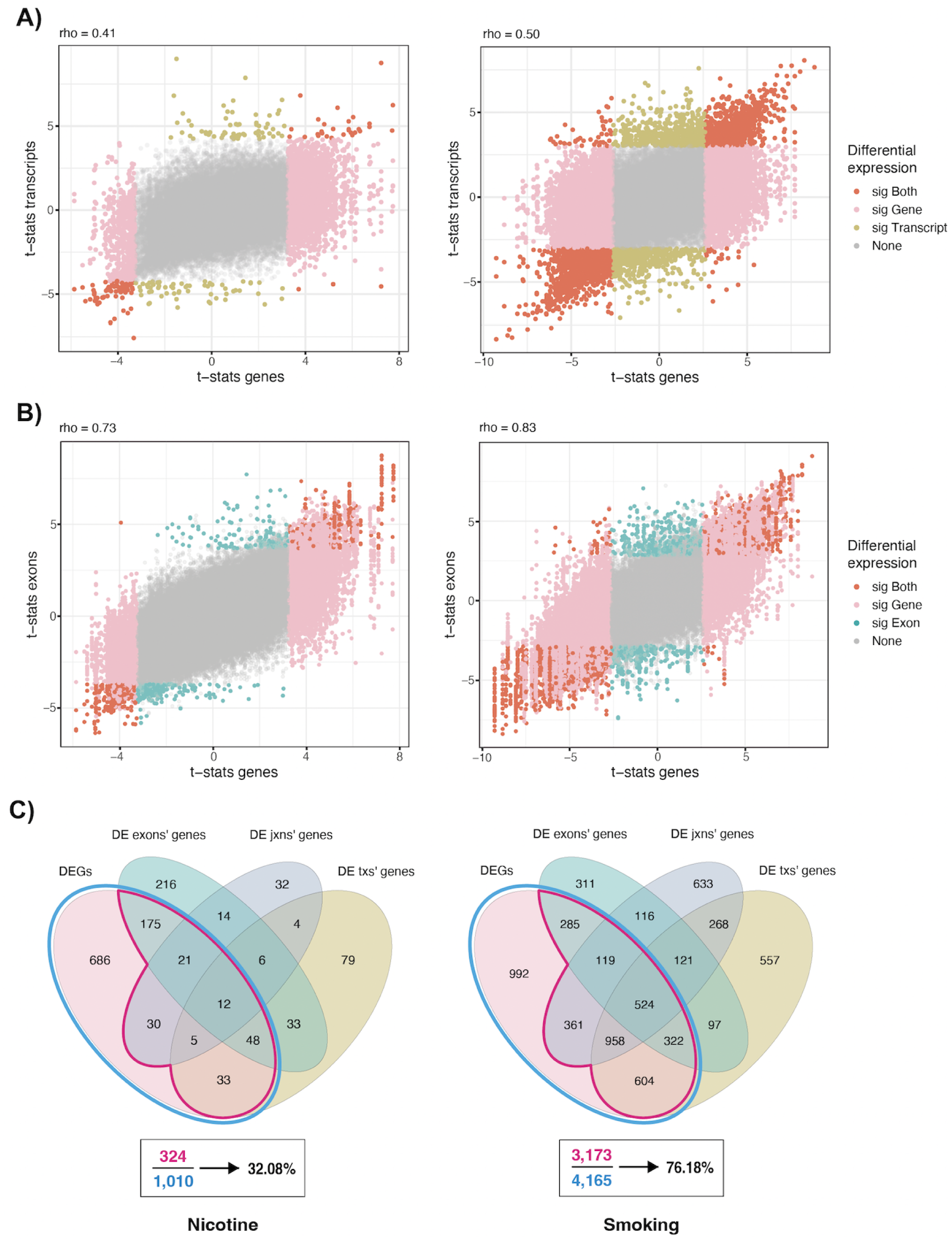
DEA results at gene, transcript, exon, and exon-exon junction levels in pup brain. Moderated *t-*statistics for differential expression of **A)** transcripts and **B)** exons in the nicotine (left) and smoking (right) experiments vs the moderated gene-level *t*-statistics in the same experiments. In dark orange DE features whose genes were also DE; in yellow and blue DE features of non-DEGs; in pink non-DE features of DEGs, and in gray non-DE features of non-DEGs. Rho corresponds to the Spearman correlation coefficient. DE transcripts and genes were defined with an FDR<5% and DE exons with FDR<5% and |logFC|>0.25. **C)** Overlap between DEGs and genes of DE transcripts (txs), exons, and exon-exon junctions (jxns) in the nicotine and smoking experiments. The percentages of DEGs with any other DE features are indicated. Related to **Fig. S17**, **Table S8**, **Table S9**, **Table S12, Table S13** and **Supplementary File 1**.

Differential exon expression (DEE) analysis identified 1,115 DE exons (FDR<0.05 and |logFC|>0.25) for nicotine exposure (**Table S10**) and 5,983 DE exons for smoking exposure (**Table S11**). Similarly to DTE, there was a strong correlation between DEE and DGE statistics (rho=0.73 and 0.83 for nicotine and smoking exposure, respectively; **Fig. 3B, Table S12, Table S13**). DE analysis at the exon-exon junction level (DJE) revealed 205 DE junctions (FDR<0.05) for nicotine exposure (**Table S14**) and 9,515 DE junctions for smoking exposure (**Table S15**). Overall, we found agreement between DE analysis results at all four expression levels, with 32% and 76.18% of the DEGs for nicotine and smoking exposure, respectively, DE at least in one other feature level (**Fig. 3C**). Functional enrichment analysis for different sets of genes based on their DE signal at the different expression levels identified synaptic vesicle and membrane components as associated with the smoking exposure (**Fig. S18A**), and overall complemented the gene-only results (**Fig. S14**). DTE, DEE, and DJE results can be used to classify DEGs based on their support at these other expression feature levels (**Fig. S19**). More fine-grained agreement for the different exposures can also be assessed to select DEGs with additional support or focus on results missed by the DGE analysis (**Fig. S20**). Further DTE, DEE, and DJE results were identified (**Supplementary File 1**).

### Molecular impact of nicotine administration and smoking exposure on adult frontal cortex and blood

To ascertain if the identified molecular impacts of nicotine and smoking exposure are specific to the developing brain, we compared those results to DGE findings for nicotine vs vehicle administration, and smoking exposure vs control in the adult brain (**Fig. 1A**). Both substances impacted differently on the gene expression in the adult brain (rho=0.03, **Fig. 4A**) but not significantly (0 DEGs at FDR<0.05, **Fig. S13A,B**), and the individual effects of each of these two substances were variable between adult vs pup brain (rho=0.01 and 0.02 for nicotine and smoking exposure, respectively, **Fig. 4B,C; Table S5**).

**Figure 4:**
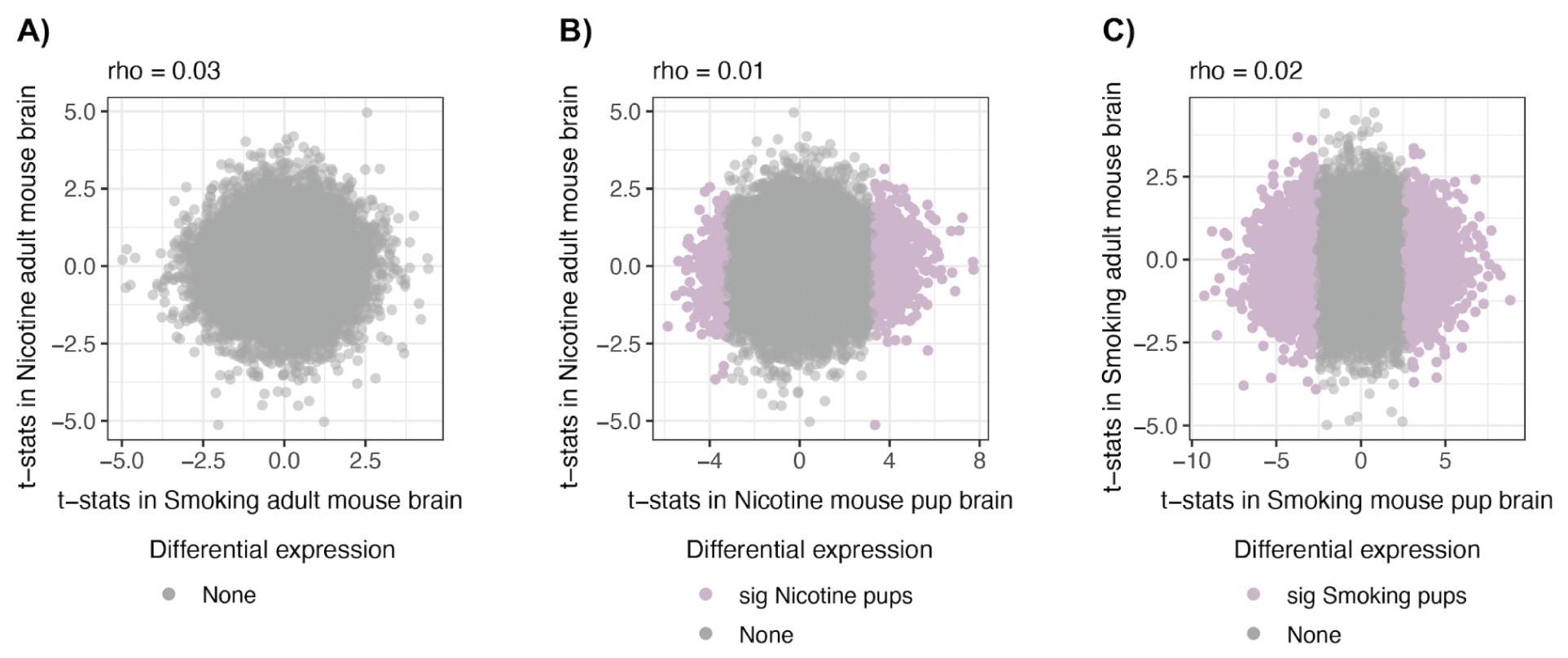
Differential gene expression signal on adult brain. Comparison of the moderated *t-*statistics of differential gene expression for **A)** nicotine administration vs smoking exposure in adult brain, and **B)** nicotine exposure and **C)** smoking exposure in adult vs pup brain. In light pink the DEGs in pup brain and in gray non-DEGs in any group; rho is the Spearman correlation coefficient. Related to **Table S5**.

We extracted RNA from blood samples of the smoking-exposed adult dams and controls (**Fig. 1A**) to evaluate if brain-level transcriptomic changes caused by smoking exposure can be read out in blood. We performed DGE for smoking exposure vs control in blood but no DEGs were found (**Fig. S13C**) and the effects of cigarette smoke in blood and brain of adults at the gene level were uncorrelated (rho=-0.01, **Fig. S21A**, **Table S5**). Nevertheless, 37 (4.8%) of the 772 genes in adult brain with nominal differences (p<0.05) for smoking exposure vs control also had nominal differences in adult blood for smoking exposure (**Fig. S21A, Table S5**). And 3% of the smoking exposure-associated DEGs in pup brain replicated in smoking-exposed adult blood (rho=-0.04, **Fig. S21B**, **Table S5**). We also identified *KCNN2*, a human gene downregulated for smoking exposure in prenatal human brain (FDR<0.1) (12), replicating in smoking-exposed mouse blood (rho=0.03, **Fig. S21C**, **Table S16**).

### Comparison of mouse transcriptomic changes with findings in human

In a previous study, the transcriptional impacts of prenatal and adult exposure to smoking on human prefrontal cortex were assessed using 33 prenatal and 207 adult, non-psychiatric postmortem brain samples, respectively. The smoking-exposed phenotype was defined by nicotine or cotinine detectability. MSDP was directly associated with differential expression of 14 genes (FDR<0.1; 16 smoking-exposed vs 17 unexposed prenatal tissue samples), whereas only 2 genes were significantly differentially expressed in adult samples (FDR<0.1; 57 active smokers vs 150 non-smokers) (12).

We used the transcriptomic results of this study to assess the replicability of our mouse differential gene expression in human. Globally, we found uncorrelated effects of smoking exposure on mouse pup and adult brain compared against prenatal and adult postmortem human brain, respectively (**Fig. 5A,B**). Nevertheless, 267 out of 4,165 (6.41%) pup brain DEGs for smoking exposure replicated in the smoking-exposed human prenatal brain (rho=-0.06, **Fig. 5A**) and 9 out of 772 (1.17%) nominally DE genes (p<0.05) in the smoking-exposed adult mouse brain replicated in the smoking-exposed human adult brain (rho=-0.01, **Fig. 5B**, **Table S16**). In particular, *NRCAM* that encodes a cell adhesion protein required for cell-cell contacts in the brain, and its mouse ortholog, were significantly downregulated in smoking-exposed human prenatal and mouse pup brain, respectively (**Fig. 5A**). *MARCO* that encodes for a pattern recognition receptor (PRR) on immune cells, as well as its ortholog in mouse, were downregulated in the smoking-exposed human adult brain (12) and in the nicotine-exposed mouse pup brain, respectively (rho=-0.03, **Fig. 5C**). Moreover, the DEGs *MPPED1* and *SDC1* in the smoking-exposed prenatal human brain replicated in the nicotine-exposed mouse pup brain (rho=-0.01, **Fig. 5D**, **Table S16**).

**Figure 5:**
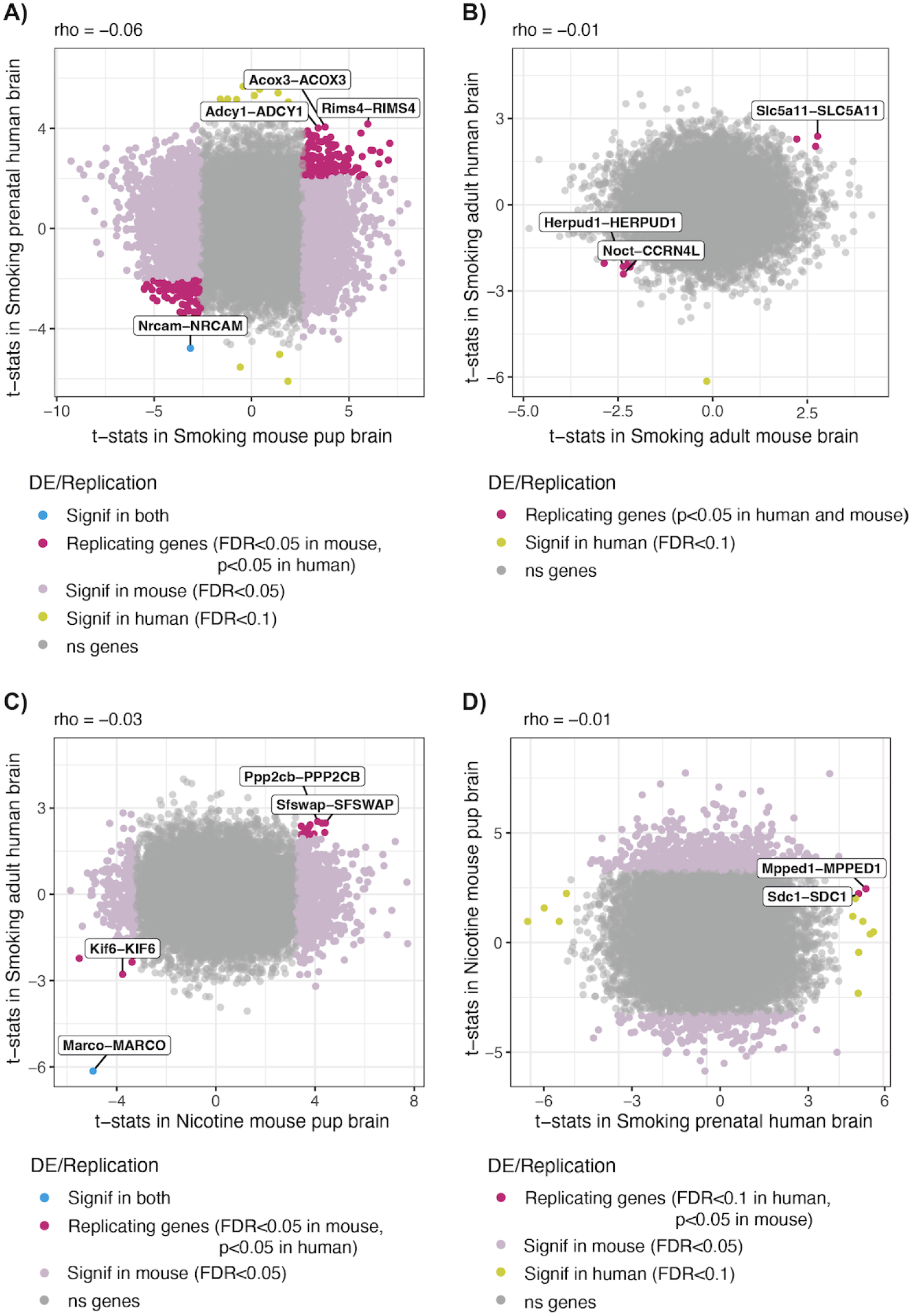
Differential gene expression signal for smoking exposure in mouse and human brain. Moderated *t*-statistics of the mouse genes for DE by (**A,B**) smoking exposure and (**C,D**) nicotine exposure in (**A,C,D**) pup and (**B**) adult mouse brain, compared against the moderated *t*-statistics of their human orthologs for smoking exposure in (**A,D**) prenatal and (**B,C**) adult human brain. In dark pink mouse (A-C) and human (D) brain genes that replicate in the other specie (with *p*-value<0.05 and same logFC sign); in light pink the genes that were DE in pup brain (FDR<0.05); in yellow the genes DE in human brain(FDR<0.1); in blue orthologous gene pairs that were DE in both species, and in gray non-DEGs in any specie. The gene pairs DE in both species, as well as the unique or the three replicating genes most significant in human are labeled with their mouse and human gene symbols. The Spearman correlation coefficient (rho) is shown above each plot. Related to **Table S16**.

Lastly, nicotine- and smoking-associated DEGs in the developing pup brain overlapped with candidate risk genes for tobacco use disorder (TUD), as identified in a genome-wide association study (GWAS) meta-analysis (25). Among the DEGs in offspring, human orthologs for *Trim35* and *Nr6a1* for nicotine exposure and *Chrna3*, *Rbm5*, *Sema3f*, and *Nfasc* for smoking exposure were associated with TUD and showed prenatal-specific expression, whereas the nicotine DEG *Vrk2* and the smoking DEGs *Drd2*, *Mtmr2*, and *Chrna3* were TUD-associated in the adult human brain. *Ip6k1* and *Cep57* were DE for both exposure experiments in pups and also associated with TUD. *Gmppb* and *P4htm* were two additional DEGs in smoking-exposed pup brain whose human orthologs were predicted to be affected in their expression by European single nucleotide polymorphisms (SNPs) TUD-associated in the human frontal cortex (**Table S17**).

## DISCUSSION

### Findings in pup and adult frontal cortex for nicotine and smoking exposure

This study interrogated transcriptomic effects of gestational smoking and nicotine exposure to the mother’s brain as well as the developing brain of the offspring in the mouse. Similar to observations in the smoking-exposed prenatal and adult human brain (12), we saw a wide signature for DE in the pup brain compared to the adult mouse brain. Likewise, reduced similarities in gene expression differences in the mature and developing brain were noted. Both findings are consistent with a stronger response to early compared to adult exposure (26,27).

Moreover, the effect of prenatal smoking exposure was more widespread than that of nicotine alone (4,165 vs 1,010 smoking-exposed and nicotine-exposed DEGs, respectively). This difference could be partly attributable to the larger number of samples used to model smoking exposure, but is also likely due to the composition of cigarette smoke, which contains >7,000 different chemicals besides nicotine (5). Therefore, although the overlap between genes affected by cigarette smoke and nicotine exposure was predictable, their effects were substantially different. Indeed, DEGs that were regulated in opposite directions in one and the other exposure reveal the differential impact on the same genes by nicotine alone and when interacting with thousands of other compounds present in the cigarette smoke. In addition, we cannot rule out differences in housing between the two experiments. Experiments were conducted in different facilities, and minor differences in standard housing, feed, or caging could contribute to differences across cohorts. Consequently, pups that were born to differently treated mice can also show experiment-dependent changes in gene expression.

Prenatal nicotine exposure significantly upregulated expression of *Foxn3* and *Arrdc3*. The former is essential for mice craniofacial development (28) and is associated with addictive substance use and compulsive behaviors in humans (29), whereas the second encodes an alpha-arrestin associated with neuroprotection in Parkinson’s disease (30) and acts as a regulator of locomotion (31), which agrees with previous results showing that PNE increases locomotor activity in mice (32,33). However, *Arrdc3*’s role in brain development remains to be explored. PNE also caused the downregulation of *Senp8*, which is involved in neural development (34), and *Coa4* which encodes a cytochrome *c* oxidase (COX) assembly factor whose downregulation may be linked to Leigh Syndrome or other related neurological disorders (35,36).

For maternal smoking during pregnancy, *Top2a* was the most significantly affected gene and was downregulated. This gene encodes for the DNA topoisomerase II alpha that regulates pluripotency and differentiation of embryonic stem cells (ESCs) (37). It has been demonstrated that maternal exposure to cigarette smoke components, such as metabolites of benzene, cause the transformation of the *Top2a* product into dangerous “molecular scissors” that fragment the genome and damage DNA in developing embryos (38). And it has been shown that the prenatal inhibition of *Top2a* causes postnatal autism-related behavioral defects in mice (39). *Tpx2* was the second most downregulated gene by prenatal smoking exposure and it plays crucial functions in the division, positioning, and fate of neural stem cells during mouse brain development (40). *AB041806* is a lncRNA gene and was the most upregulated after smoking exposure in pups; it is expressed in the CNS but the existence of its encoded protein has not been experimentally proven and its involvement in brain function is not known yet (41,42). The second most upregulated gene for prenatal smoking exposure was *Mt2*, which encodes a metallothionein (MT), a metal-binding protein that acts as an antioxidant and whose expression is known to be induced in the CNS as a response to brain damage (43). In fact, previous studies have identified *Mt2* as significantly upregulated in astrocytes after cerebral ischemic damage in mice (44) and following induced seizure attack in rats (45). In human MT genes have been found upregulated in astrocytes of patients with Alzheimer’s and Parkinson’s diseases (43,46,47).

Notably, genes that were upregulated after both PNE and MSDP play relevant roles in the cellular distribution and formation of protein complexes composed of several Sm proteins and the survival motor neuron (SMN) protein, such as *Strap*, *Snrpd3*, and *Snrpd*. The SMN-Sm complex is essential for spliceosomal small nuclear ribonucleoproteins (snRNPs) assembly in the cytoplasm for pre-mRNA splicing events and the highest levels of activity of this complex occur during embryonic and early postnatal development of the CNS (48). Other DEGs such as *Nsg1*, *Clstn1*, and *Rab4a* that were also upregulated in both experiments act in postsynaptic endosomes, which contribute to neural development regulation (49).

Genes uniquely upregulated by nicotine and not by smoking exposure were also identified, such as *Gsk3a*, *Ppp2r2b*, and *Ppp1cc* that are implicated in dopaminergic synapses, *Gnai3* and *Gnaq* that are related to synaptic long-term depression (LTD), as well as *Ppp2cb* which is involved in both pathways. These results are in alignment with previous findings reporting the nicotine interference in the dopamine neurotransmitter system development (11,50,51) and the nicotinic activity in LTD induction in rat and mouse brains (52–54). Besides, *Cplx2*, *Ykt6*, and *Cplx3* whose products participate in the SNARE complex were specifically upregulated by smoking but not by nicotine exposure, which could have relevant neurocognitive and behavioral implications, in support of the observed relationship between deficits in the SNARE protein SNAP-25 and maternal smoking with Attention Deficit Hyperactivity Disorder (ADHD) (33,55,56). Accordingly, the gestational exposure to smoking was associated with the upregulation of transcripts whose products act in the SNARE complex and are involved in vesicular transport, as well as transcripts of genes involved in Parkinson’s, Huntington’s, and prion-related diseases, such as *Casp3*, *Psma6*, and *Nduvf1*, placing MSDP as a potential not yet fully addressed environmental factor linked to the susceptibility or development of neurodegenerative disorders in offspring (57–59).

Together, the differential expression of these genes demonstrates a variable impact of cigarette smoke and nicotine on brain development and introduces potential long-term effects on the offspring related to neurodegenerative, neurodevelopmental, and substance use disorders. In the future, it will be informative to monitor behavioral and cognitive traits of the exposed newborn to gain insights into the postnatal effects related to prenatal nicotine and smoking exposure, as well as their underlying molecular processes including epigenetic modifications that may mediate the effect of nicotine and smoking exposure on gene expression.

### Blood vs brain molecular changes by smoking exposure in mice

Numerous epidemiological and toxicological studies have analyzed smoking effects in blood as an approach to establish brain effects. These studies either rely on the hypothesis that smoking components affect cardiovascular and brain health through the same responsive mechanisms, or that brain perturbations are, at least in part, a consequence of cardiovascular effects via circulation of pro-inflammatory mediators or ultrafine particulate matter that can reach the brain (60–62). Here we compared transcriptomic alterations caused by cigarette smoke exposure in adult brain and blood and found uncorrelated effects. Concordant with a previous investigation (63), this suggests that the study of cigarette smoke impact on brain cannot be addressed merely by the examination of blood samples.

Nonetheless, we found *Dusp14* downregulated in both smoking-exposed adult brain and blood (*p*-value<0.05 in both tissues). This gene regulates inflammation and oxidative stress and has been found downregulated in the infarcted area of mice after ischemic stroke (64), possibly linking tobacco exposure effects in brain and blood given that smoking is a well-recognized risk factor for stroke (62,65). *Syt13* was another gene downregulated in both tissues (*p*-value<0.05) with a role in neurotransmitter secretion by synaptic vesicles (66) but also recently characterized in human as a biomarker in lung adenocarcinoma (67), consistent with smoking exposure. A third adult brain gene replicating in blood was *Arhgef25*, also downregulated and whose human ortholog is expressed in brain vasculature (66). The upregulated DEG *Pde3b* in smoking-exposed pup brain also replicated in blood. Its expression is known to be increased after ischemic insult in the mouse brain (68) and accordingly its deletion/inhibition confers protection from ischemia/reperfusion (I/R) injury in mouse heart (69). The following most significant pup DEGs for smoking exposure replicating in blood were *Arhgap28* and *Slc39a6*, both downregulated and which need to be more widely studied in order to determine the relationship between their cigarette smoke effects on brain and blood. The downregulated DEG *KCNN2* in the smoking-exposed human prenatal brain (12) replicated in mouse blood. This gene is expressed in mouse brain and heart and several of its polymorphisms have been associated with cardiac tachyarrhythmias in human (70) and neurodevelopmental movement disorders and locomotor deficits in both humans and rodents (71,72). Moreover, its expression in brain is relevant for alcohol, nicotine, and drug addiction (73).

### Coincident molecular changes by smoking and nicotine exposure in mouse and human brain

Lastly, we explored to what extent our mouse results can be extrapolated to human using DGE results for smoking exposure in prenatal and adult human brain (12). An advantage of using mice to study the effects of prenatal and adult drug exposure is the ability to control experimental conditions that circumvents the confounding implications of human factors commonly coincident with drug use that also have impact on the brain, such as poor prenatal care and exposure to other substances (4), which makes it difficult to identify specific substance effects with certainty. However, the different gestation periods, routes of administration, pharmacokinetics, and correlation between transcriptomes of pups from the same litter, are some of the limitations of modeling these processes in animal models that hinder translatability to humans (4). In fact, our results indicate that the impacts of smoking exposure on gene expression in mouse and human brain are variable, which besides being explained by the inherent biological differences between species and experimental challenges modeling these processes, can be conceivable in terms of variations in the RNA-seq data processing steps and in the formal DGE analysis.

Nevertheless, DGE signal replicated between smoking/nicotine-exposed mouse brain and smoking-exposed human brain (12). The human gene *NRCAM* and its mouse ortholog were downregulated in the developing brain after cigarette smoke exposure (12). This gene encodes a neuronal cell adhesion protein with essential roles in axon growth and guidance and the formation of neural circuitry during brain development (74–80). *Nrcam*-null or deficient mice present autism-related behavioral and phenotypic alterations (75,77) and changes in its expression are associated with psychiatric disorders and drug addiction (74). *MARCO* was a downregulated gene in the smoking-exposed human adult brain (12) whose mouse ortholog was also downregulated in the nicotine-exposed pup brain, defining a gene expression change that is preserved regardless of species, age, and experiment setup. The product of this gene is a macrophage receptor with collagenous structure expressed in microglia involved in neuroinflammatory responses in neurodegenerative diseases (81,82). Its unknown involvement in neurodevelopment matches with its age-independent differential expression but it was not surprising to find it DE as it has been demonstrated that cigarette smoke exposure significantly decreases the expression of this gene in macrophages, which in turn leads to decreased pathogen clearance (83,84). Therefore, our results suggest nicotine and smoking can compromise brain immune function, as has been previously proposed (59,62).

Finally, finding DEGs in pup brain for both nicotine and smoking exposure, whose human orthologs are TUD-associated (25) with a matching brain region or developmental stage-specific expression, further suggests that MSDP and PNE can increase the likelihood of experimenting with drugs later in life, as has been extensively reported (85–92).

In summary, the present study revealed nicotine-specific and broader cigarette smoke transcriptomic effects on mouse brain development. The gene-level results were consistent and complemented with evidence at the transcript, exon, and exon-exon junction levels, finding DEGs and genes with other DE features with clear involvement in neurodevelopmental and behavioral processes. Also demonstrated were the variable effects of nicotine and cigarette smoke on the pup and adult mouse brain, as well as the non-extrapolable impact of tobacco smoke from mouse blood to brain, though, as presented, some genes subject to additional research could serve as biomarkers for smoking in these two tissues. Finally, these findings were supported by several human genes TUD-associated or affected by smoking in the prenatal and adult human prefrontal cortex that were also DE in the nicotine- and smoking-exposed pup brain. In conclusion, new insights into the genes and pathways implicated in the deleterious developmental effects of nicotine and cigarette smoke exposures during gestation were found and valuable data useful for ongoing research regarding the effects of MSDP and PNE were generated and publicly shared.

## ACKNOWLEDGEMENTS

We thank the Joint High Performance Computing Exchange (JHPCE) for providing computing resources for these analyses. We thank Louise A. Huuki-Myers (Lieber Institute for Brain Development; LIBD) for guidance in data visualizations. We acknowledge Brianna K. Barry (JHU) for generating RNA-seq libraries.

## Funding

This project was supported by National Institutes of Health grants R01EY033765 (JTH), R01EY031594 (JTH), R01EY035805 (JTH), R01DA042090 (DBH, AEJ), and R01MH105592 (KM) and the Lieber Institute for Brain Development (LIBD).

## Code availability

Code for data analysis of this project can be found in the GitHub repository https://github.com/LieberInstitute/smoking-nicotine-mouse (93).

## Data availability

Unfiltered and normalized expression data generated and used in this project is available through the *smokingMouse* (94) Bioconductor data package, which also includes sample and feature-level data and the results from the DEA on human frontal cortex (12). Raw bulk RNA-seq files are available from the NCBI Sequence Read Archive (BioProject PRJNA1175674).

## Abbreviations

DE: differentially expressed or differential expression.
DEA: differential expression analysis.
DEE: differential exon expression
DEGs: differentially expressed genes
DGE: differential gene expression.
DJE: differential exon-exon junction expression.
DTE: differential transcript expression.
FDR: false discovery rate.
GO: Gene Ontology.
GWAS: genome-wide association study.
KEGG: Kyoto Encyclopedia of Genes and Genomes.
MSDP: maternal smoking during pregnancy.
P0: postnatal day 0.
PNE: prenatal nicotine exposure.
SNP: single nucleotide polymorphisms.
TUD: tobacco use disorder.

## Author contributions

Conceptualization: AEJ, KM Methodology: DGP, LCT Investigation: MC, KM Software: DGP, NJE, LCT Formal Analysis: DGP

Data Curation: NJE Writing-original draft: DGP

Writing-review and editing: GP, KRM, KM, LCT Visualization: DGP

Supervision: JTH, LCT, KM Project administration: KM

Funding Acquisition: AEJ, DBH, KM

## CONFLICT OF INTEREST

Andrew E. Jaffe (AEJ) is currently a full-time employee at Neumora Therapeutics. AEJ’s current work is unrelated to the contents of this manuscript, and his contributions to this manuscript were made while previously employed at LIBD. No other authors have financial relationships with commercial interests, and the authors declare no competing interests.

## SUPPLEMENTARY MATERIAL

### Supplementary Materials and Methods

#### Animals

Wild type male and female mice (C57BL/6J; stock # 000664, Jackson Laboratories) were purchased and used for timed breeding. 6 week old female mice were paired with male mice. Copulation plugs were checked daily and male mice were removed upon identification of plugs. Female mice were monitored for pregnancy, and separated upon pregnancy confirmation. Pups were euthanized by decapitation on the first day following birth, e.g. postnatal day 0 (P0). Pregnant dams were euthanized by decapitation and trunk blood was collected into a heparinized tube. Brains were rapidly extracted from the skull and the frontal cortex was dissected from the brain over wet ice on a steel block using a scalpel. Frontal cortical tissue was snap frozen in chilled 2-methylbutane. Samples were transferred to tubes and placed on dry ice and stored at -80°C until further processing for RNA extraction. All experiments and procedures were approved by the Johns Hopkins Animal Care and Use Committee and in accordance with the Guide for the Care and Use of Laboratory Animals.

#### Nicotine administration

Free-base(-)-nicotine (Sigma) was dissolved in normal saline. Nicotine (1.5mg/kg) or vehicle (saline) was administered to female dams (2X/daily - 8AM and 4PM). Administration started the week before mice were paired and continued until E17.

#### Smoking exposure

Pregnant dams were placed into a smoking chamber for 5 hours/day, 5 days/week starting one week before mice were paired for breeding and until the time of delivery. This chamber contains a smoking machine (Model TE-10, Teague Enterprises, Davis, CA) that burns 5 cigarettes (2R4F reference cigarettes (2.45 mg nicotine/cigarette; Tobacco Research Institute, University of Ky) at a time, taking 2 second duration puffs at a flow rate of 1.05 l/min, to provide a standard puff of 35 cm^3^, providing a total of 8 puffs per minute. The machine is adjusted to produce side stream (89%) and mainstream smoke (11%). The chamber atmosphere is monitored to maintain total suspended particulate at 90 mg/m^3^, and carbon monoxide at 350 ppm. Control pregnant dams were kept in a filtered air environment.

#### Tissue processing and RNA isolation and sequencing

Total RNA was extracted from samples using Trizol followed by purification with an RNeasy Micro kit (Qiagen). Paired-end strand-specific sequencing libraries were prepared and sequenced by Macrogen from 1ug total RNA using the TruSeq Stranded mRNA kit with ERCC

Spike in. Libraries were sequenced on an Illumina NovaSeq6000 S4, 150bp paired end. Output was targeted at 60M total reads (R1 30M and R2 30M) million 150-bp paired-end reads.

#### RNA-seq data processing

### Expression quantification

Quality assessment of the sequencing reads and expression quantification at gene, exon, transcript, and exon-exon junction levels were performed running the RNA-seq processing pipeline *SPEAQeasy* version 6c1dab0 (13), using default settings, which involved alignment of reads to the Mus musculus genome from GENCODE M25 (95,96). As part of *SPEAQeasy, featureCounts* (19) was used for gene and exon quantification, using the `-O` argument for exon features in order to assign reads to all the overlapping exons. This setting has the drawback that it can inflate exon read counts. *RegTools* (97) was used for junction quantification and *Kallisto* (98) performed the pseudoalignment of the reads to the transcriptome.

### Count normalization

Raw counts were normalized by sample library size calculating normalization factors with `calcNormFactors()`function from *edgeR* v3.43.7 (99) using the trimmed mean of M-values (TMM) method (100) for genes and exons, and the TMM with singleton pairing (TMMwsp) method for junctions. After rescaling library sizes, counts were transformed into counts per million (CPM) in a logarithmic scale with the *edgeR* `cpm()`function. For transcripts, transcripts per million (TPM) were log2-transformed after adding a 0.5 scaling factor (**Fig. S1**).

### Feature filtering

Lowly-expressed genes, exons, and exon-exon junctions were filtered based on their counts across samples using `filterByExpr()`from *edgeR* v3.43.7 (99), which retains features that have a minimum of 15 reads across all samples and at least 10 reads in *n* or more samples, where *n* is 70% the size of the smallest sample group. Transcripts were filtered by defining a mean TPM expression cutoff of 0.28 with `expression_cutoff()`from *jaffelab* v0.99.32 (101) (**Fig. S1**).

#### RNA-seq data analysis

### Exploratory data analysis

Quality control (QC) metrics of the samples were direct outputs of the *SPEAQeasy* pipeline (13) and additional ones were computed using `addPerCellQC()`from *scuttle* v1.9.4 (102) that can operate on counts at sample-level; all these metrics were calculated from raw counts of genes before normalization and filtering. These metrics were examined and compared separately for brain and blood samples, detecting large differences in the proportions of reads that mapped to mitochondrial and ribosomal genes in adult and pup brain samples (**Fig. S2**), which led to age explaining a high percentage of gene expression variance in brain samples (**Fig. S3A**) and directed further separation of brain samples by age for downstream analyses.

Poor-quality samples were defined as those presenting lower outlier values for library size or number of detected genes, or higher outlier values for the percentage of either mitochondrial or ribosomal read counts. Values were considered outliers if they were 3 median-absolute-deviations away from the median, as defined by `isOutlier()` from *scater* v1.29.1 (102) (**Fig. S4**). After removing those samples, sources of gene, transcript, exon, and exon-exon junction expression variation in the samples were explored through dimensionality reduction analyses. Principal Component Analysis (PCA) revealed big differences in gene, transcript, and exon expression of brain samples from adult mice that were part of the nicotine and smoking exposure experiments, including both exposed and control samples each (**Fig. S5**, **Fig. S7A**), and exhibited 3 poor-quality samples that appeared isolated from the rest in PC plots (**Fig. S5**); these were manually filtered out. Similarly, three segregated poor-quality samples from pup brain were identified in PC plots, and sex appeared as a major driver of gene and exon expression variability (**Fig. S6**, **Fig. S7B**). In blood samples, pregnancy state slightly contributed to transcriptomic differences (**Fig. S3B**). Posterior to QC-based and manual sample filtering, 23 blood samples, 39 adult brain samples, and 130 pup brain samples were kept for downstream analyses (**Table S2**). Multidimensional scaling (MDS) analysis corroborated PCA results in the filtered adult and pup brain samples (**Fig. S8**, **Fig. S9**, **Fig. S10**).

In order to guide the selection of sample variables to include in the models for differential expression analysis, for each gene the percentage of expression variance explained by each individual explanatory variable was computed with `getVarianceExplained()`from *scater* v1.29.1 (102) (**Fig. S11**), as well as the fractions of variance explained (FVE) by each variable accounting for the joint contribution of all of them with `fitExtractVarPartModel()`from *variancePartition* v1.32.2 (103) (**Fig. S12**). In that way, QC metrics and biological variables such as pregnancy and sex, in adult and pup samples, respectively, were identified as major contributors to changes in the gene expression profiles. Pairs of highly correlated variables were recognized running a Canonical Correlation Analysis (CCA) with the *variancePartition* function `canCorPairs()`and only the one variable with the highest median FVE from each pair was added to the models (**Fig. S12**).

### Differential expression analyses

Differentially expressed features for nicotine vs vehicle exposure/administration, and smoking exposure vs control were identified in pup brain, and adult brain and blood by defining the following models:

- For smoking exposure vs control in adult blood, and both nicotine vs vehicle administration and smoking exposure vs control in adult brain (analysis only at the gene level):

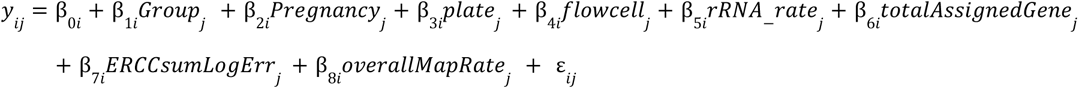

- For nicotine vs vehicle exposure, and smoking exposure vs control in pup brain (analysis at the four levels of expression features):

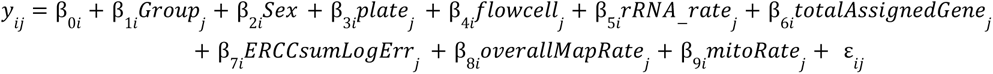

Where *y*_*ij*_ denotes the expression of the *i*th feature in the *j*th sample, modeled by the selected covariates (see **Table S18** for their description) plus an error term ε.

We applied an empirical Bayes analysis pipeline with *limma* v3.57.6 (14) for differential expression analysis. For gene, transcript, and exon counts, `voom()`was used as a first step to estimate inverse variance weights for each expression observation based on the mean-variance trend of the data to adjust for count heteroscedasticity (**Fig. S13**). This function renormalized raw counts into log-CPM using the previously computed normalization factors and library sizes for the non-filtered datasets. The log-normalized counts and their associated weights, as well as log-TPM of transcripts, were subsequently entered into `lmFit()`to fit a linear model by weighted least squares for each feature and estimate the model coefficients. Then `eBayes()` was used to moderate the residual sample standard deviations of the transcriptomic features through an empirical Bayes model. Finally *p*-values of the resulting moderated *t*-statistics were adjusted for multiple testing with the Benjamini and Hochberg’s (BH) method (104) to control the FDR using `topTable()`(**Fig. S13**). Only genes, transcripts, and exon-exon junctions with an FDR<0.05, as well as exons with an FDR<0.05 and |log2FC|>0.25, were considered differentially expressed.

Replication of mouse brain DE results in mouse blood or human brain was defined with an FDR<0.05 for pup brain/*p*-value<0.05 for adult brain, a *p*-value<0.05 in blood/human, and the same regulation directionality in both tissues/species. Replication of human brain DE in mouse blood/brain was defined with an FDR<0.1 in human, *p*-value<0.05 in mouse, and same regulation directionality in both. Note however that when contrasting mice and human, results of gene pairs (mouse-human orthologs), but not individual genes, are compared.

### Differential gene expression visualization

The z-scores of the log-normalized counts of DEGs were computed to visualize their expression patterns in heat maps, agglomerating genes and samples by expression through complete-linkage hierarchical clustering using an euclidean distance measure.

### Novel junction gene annotation

The nearest (overlapping) neighbor and closest downstream and upstream (non-overlapping) genes of DE novel junctions were found using the functions nearest(), precede() and follow(), respectively, from *IRanges* v2.36.0 (105).

### Software

*ggplot2* v3.4.4 (106), *R* version 4.3.0 (107), and *Bioconductor* version 3.18 (108) were used to perform all the analyses and visualize the results.

## Supplementary Tables

Supplementary Table 1: Study design. Number of samples from each pair of sample-level variables. See Table S18 for sample variable description.

Supplementary Table 2: Samples used for downstream analyses. Number of samples from each pair of sample-level variables after sample filtering based on QC metrics and PCA plots. See Table S18 for sample variable description.

Supplementary Table 3: DEGs in the nicotine-exposed pup brain. Metadata of DEGs in the nicotine pup brain and their logFC, moderated *t*-stats, *p*-value, and adjusted *p*-value for DE. The statistics were computed with topTable() from *limma*; see its documentation for the definition of the variable names. Related to Fig. 2A.

Supplementary Table 4: DEGs in the smoking-exposed pup brain. Same as in Table S3 but for DEGs in the smoking pup brain. Related to Fig. 2B.

Supplementary Table 5: Differential gene expression results for the complete gene dataset. Gene-level metadata and the logFC, moderated *t*-stats, *p*-value, and adjusted *p*-value for DE of each gene in the 5 experimental groups: nicotine vs vehicle exposure in pup brain, smoking exposure vs control in pup brain, nicotine vs vehicle administration in adult brain, smoking exposure vs control in adult brain, and smoking exposure vs control in adult blood. Also included are the replication results of the genes in mouse blood. The statistics were computed with topTable() from *limma*; see its documentation for the definition of the variable names. Related to Fig. 2C, Fig. 4 and Fig. S21A-B.

Supplementary **Table 6**: Differentially expressed transcripts in the nicotine-exposed pup brain. Metadata of DE transcripts in the nicotine pup brain and their logFC, moderated *t*-stats, *p*-value, and adjusted *p*-value. The statistics were computed with `topTable()` from *limma*; see its documentation for the definition of the variable names.

Supplementary **Table 7**: Differentially expressed transcripts in the smoking-exposed pup brain.

Same as in Table S6 but for DE transcripts in the smoking pup brain.

Supplementary **Table 8**: Differential expression of transcripts vs genes for the nicotine experiment in pup brain. DE statistics (logFC, moderated *t*-stats, *p*-value, and adjusted *p*-value) for transcripts and their respective genes for nicotine exposure in pup brain, and if transcripts and genes were both or solely DE. Only transcripts of genes present in the gene dataset are shown. The statistics were computed with `topTable()` from *limma*; see its documentation for the definition of the variable names. Related to Fig. 3A.

Supplementary **Table 9**: Differential expression of transcripts vs genes for the smoking experiment in pup brain. Same as in Table S8 but for the smoking experiment. Related to Fig. 3A.

Supplementary **Table 10**: Differentially expressed exons in the nicotine-exposed pup brain. Metadata of DE exons in the nicotine pup brain and their logFC, moderated *t*-stats, *p*-value, and adjusted *p*-value. The statistics were computed with `topTable()` from *limma*; see its documentation for the definition of the variable names.

Supplementary **Table 11**: Differentially expressed exons in the smoking-exposed pup brain. Same as in Table S10 but for DE exons in the smoking pup brain.

Supplementary **Table 12**: Differential expression of exons vs genes for the nicotine experiment in pup brain. DE statistics (logFC, moderated *t*-stats, *p*-value, and adjusted *p*-value) for exons and their respective genes for nicotine exposure in pup brain, as well as if exons and genes were both DE or not. Only exons of genes present in the gene dataset are shown. The statistics were computed with `topTable()` from *limma*; see its documentation for the definition of the variable names. Related to Fig. 3B.

Supplementary **Table 13**: Differential expression of exons vs genes for the smoking experiment in pup brain. Same as in Table S12 but for the smoking experiment. Related to Fig. 3B.

Supplementary **Table 14**: Differentially expressed exon-exon junctions in the nicotine-exposed pup brain. Metadata of DE exon-exon junctions in the nicotine pup brain, including for each:

○ if both the donor and acceptor sites together are known and annotated in GENCODE M25 (`inGencode` variable);

○ if the donor and acceptor sites are individually annotated in GENCODE M25 (`inGencodeStart` and `inGencodeEnd` variables, respectively);

○ the junction class: *Novel* (if both start and end sites are unknown, also known as fully novel junctions), *InGen* (already annotated in GENCODE M25), *AltStartEnd* (if it has only one known site), or *ExonSkip* (with sites from non-successive exons, both known individually but not together), and

○ if they are fusion junctions, meaning that they connect exons from different genes (`isFusion` variable).

Their logFC, moderated *t*-stats, *p*-value, and adjusted *p*-value are provided. These statistics were computed with `topTable()`from *limma*; see its documentation for the definition of the variable names.

Supplementary **Table 15**: Differentially expressed exon-exon junctions in the smoking-exposed pup brain. Same as in Table S14 but for DE exon-exon junctions in the smoking pup brain.

Supplementary **Table 16**: Differential gene expression results for gene pairs of mouse-human orthologs. The logFC, moderated *t*-stats, *p*-value, and adjusted *p*-value of the human gene for smoking exposure in the prenatal and adult human brain, and of the corresponding mouse orthologous gene for the 5 experimental mice groups (as in Table S5), are presented. Only mouse genes with human ortholog(s) present in the human dataset from (12) are considered. The DE statistics were computed with `topTable()` from *limma*; see its documentation for the definition of the variable names. Related to Fig. 5 and Fig. S21C.

Supplementary **Table 17**: Mouse DEGs in pup brain with human orthologs TUD-associated. Pup brain DEGs for the nicotine and smoking exposure with human orthologs that were the nearest genes of genome-wide significant (GWS) lead SNPs in loci associated with TUD, obtained from a multi-ancestral GWAS meta-analysis of TUD cases and controls in individuals from 8 cohorts (including UKBB), with European (EUR), African American (AA), and Latin American (LA) ancestry (*TUD-multi+UKBB* dataset), and from a within-ancestry GWAS meta-analysis in EUR individuals from 5 cohorts, including UKBB data (*TUD-EUR+UKBB* dataset). As well as human genes significantly associated with TUD in EUR individuals (*TUD-EUR-MAGMA* dataset); neurobiologically relevant target human genes associated with TUD (*TUD-EUR-H-MAGMA* dataset), especially expressed in prenatal (*TUD-EUR-H-MAGMA-prenatal* dataset) and adult brain (*TUD-EUR-H-MAGMA-adult* dataset); TUD-associated human genes whose expression is predicted to be affected by EUR-SNPs across multiple brain regions (*TUD-EUR-S-MultiXcan* dataset) and in specific brain regions (*TUD-EUR-S-PrediXcan* dataset), including the frontal cortex (*TUD-EUR-S-PrediXcan-FC* dataset). See more details of these TUD-associated human genes in the original publication (25).

Supplementary **Table 18**: Dictionary of sample variables. Description of the sample variables used throughout this project.

Supplementary **Table 19**: Associated genes of fully novel DE exon-exon junctions in pup brain. Nearest, following, and preceding genes of the fully novel DE exon-exon junctions without assigned gene for the nicotine and smoking exposure in pup brain.

## Supplementary Figures

**Supplementary Figure 1:**
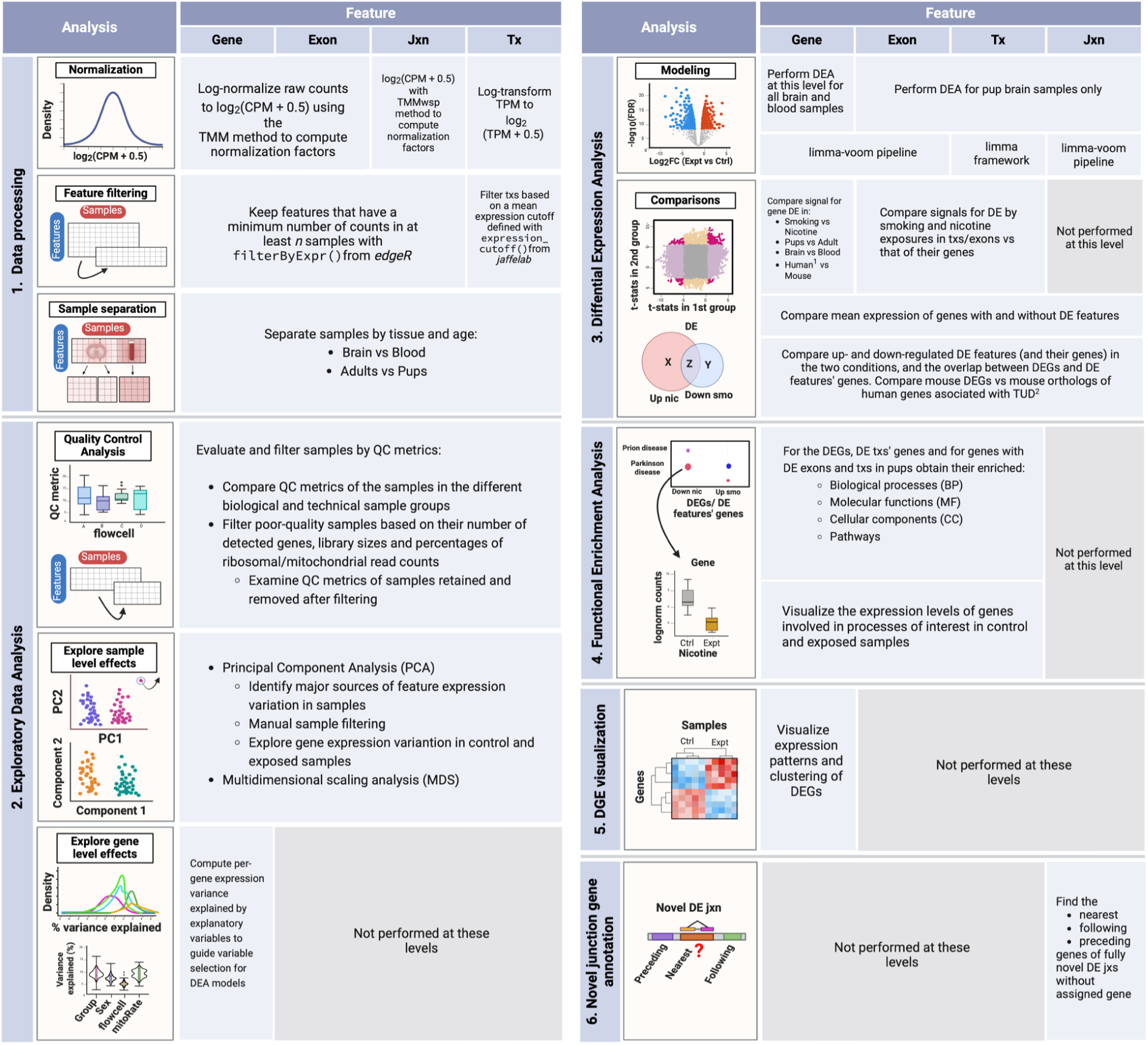
**Summary of analysis steps across gene expression feature levels**. **1. Data processing**: counts of genes, exons, and exon-exon junctions were normalized to CPM and log2-transformed; transcript expression values were only log2-transformed since they were already in TPM. Lowly-expressed features were removed using the indicated functions and samples were separated by tissue and age in order to create subsets of the data for downstream analyses. **2. Exploratory Data Analysis (EDA)**: QC metrics of the samples were examined and used to filter the poor quality ones. Sample level effects were explored through dimensionality reduction methods and segregated samples in PCA plots were removed from the datasets. Gene level effects were evaluated with analyses of variance partition. **3. Differential Expression Analysis (DEA)**: with the relevant variables identified in the previous steps, the DEA was performed at the gene level for nicotine and smoking exposure in adult and pup brain samples, and for smoking exposure in adult blood samples; DEA at the rest of the levels was performed for both exposures in pup brain only. DE signals of the genes in the different conditions, ages, tissues, and species (^1^ using human results from Semick et al., 2020) were contrasted, as well as the DE signals of exons and transcripts vs those of their genes. Mean expression of DEGs and non-DEGs genes with and without DE features was also analyzed. Then, all resultant DEGs and DE features (and their genes) were compared by direction of regulation (up or down) between and within exposures (nicotine/smoking); mouse DEGs were also compared against ^2^human genes associated with TUD from Toikumo et al., 2023. **4. Functional Enrichment Analysis**: GO & KEGG terms significantly enriched in the clusters of DEGs and genes of DE transcripts and exons were obtained. **5. DGE visualization**: the log2-normalized expression of DEGs was represented in heat maps in order to distinguish the groups of up- and down-regulated genes. **6. Novel junction gene annotation**: for uncharacterized DE junctions with no annotated gene, their nearest, preceding, and following genes were determined. See **Supplementary Materials and Methods** for complete details. **Abbreviations:** Jxn: junction; Tx(s): transcript(s); CPM: counts per million; TPM: transcripts per million; TMM: Trimmed Mean of M-Values; TMMwsp: TMM with singleton pairing; QC: quality control; PC: principal component; DEA: differential expression analysis; DE: differential expression/differentially expressed; FC: fold-change; FDR: false discovery rate; DEGs: differentially expressed genes; TUD: tobacco use disorder; DGE: differential gene expression.

**Supplementary Figure 2:**
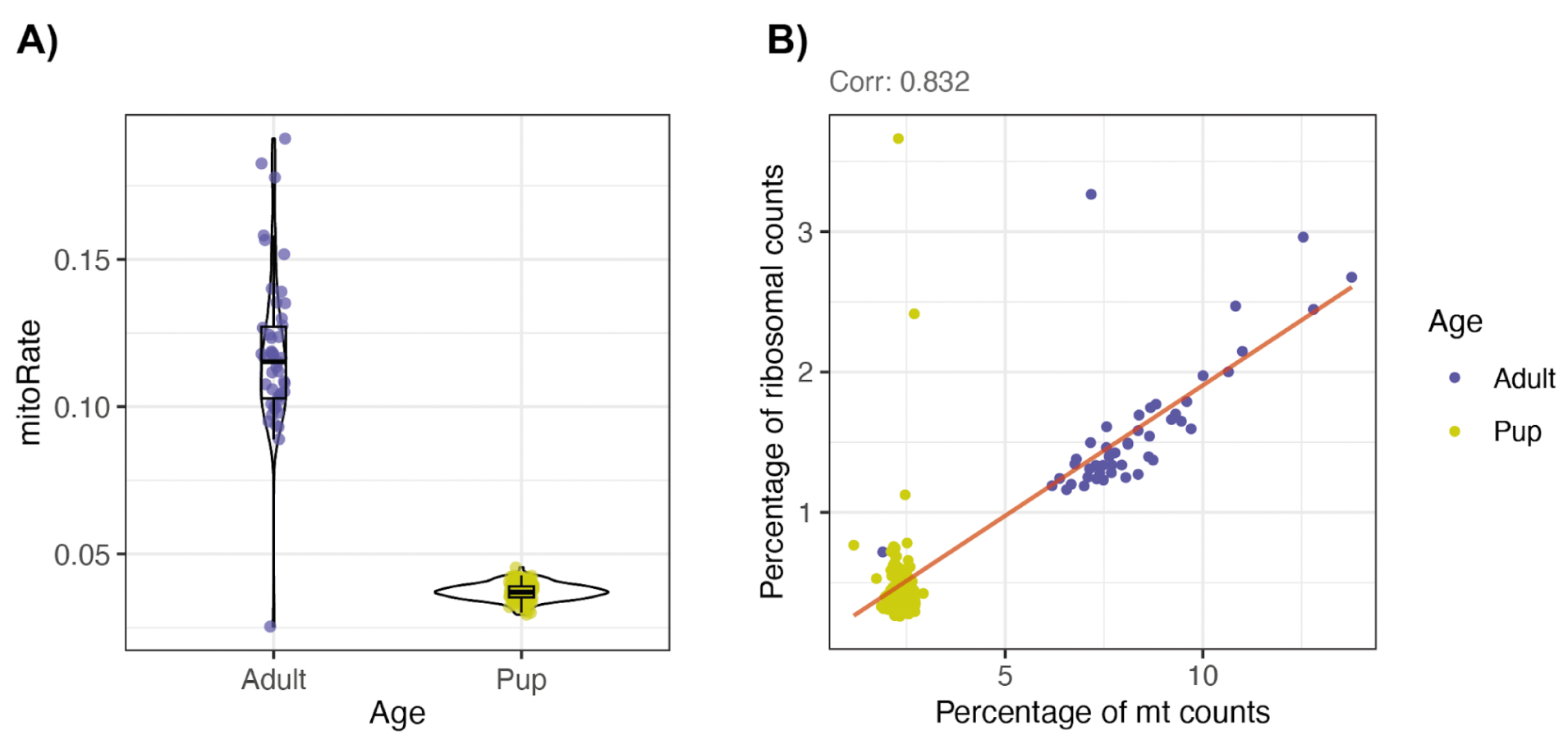
**Quality control metrics of adult and pup brain samples**. **A)** The decimal fraction of reads that mapped to the mitochondrial chromosome of those that mapped at all in adult and pup brain samples. **B)** Percentage of sample counts from reads that were assigned to mitochondrial (mt) genes vs those that mapped to ribosomal genes, for each brain sample. Pearson correlation coefficient between these two QC metrics is shown above and the fitted linear regression line is shown in red.

**Supplementary Figure 3:**
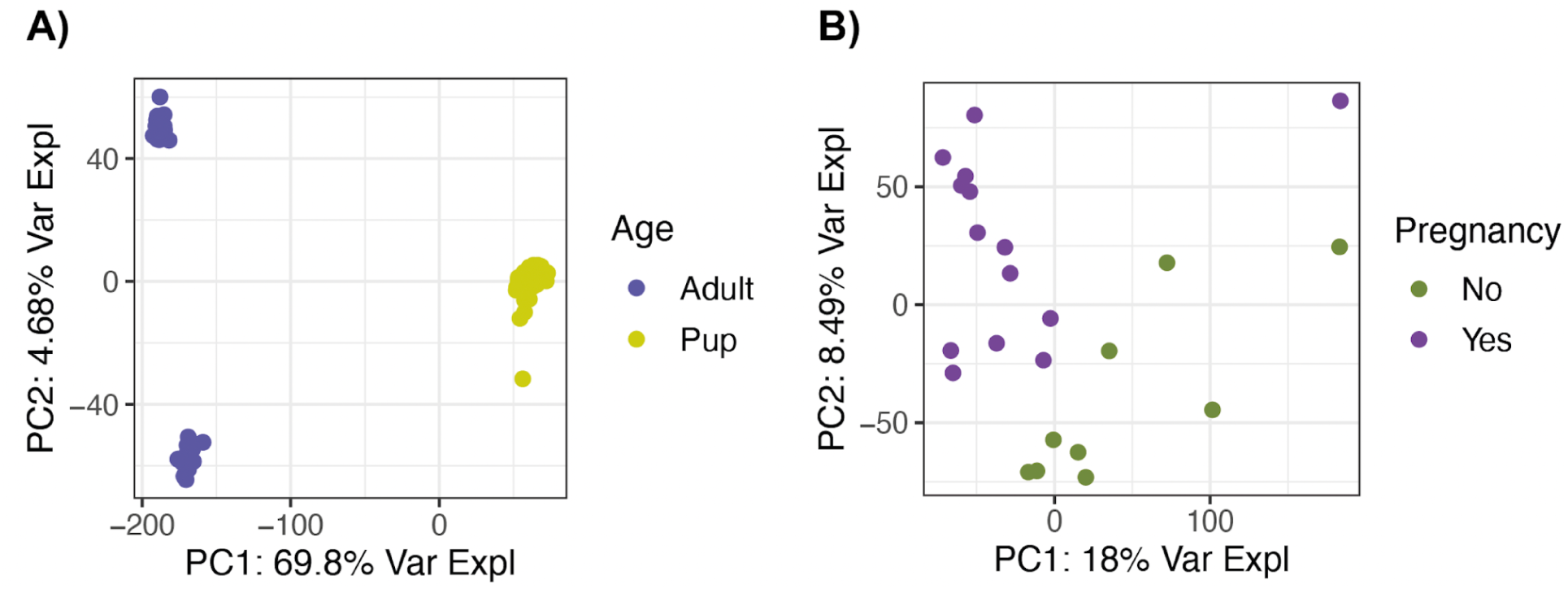
Principal Component Analysis in brain and blood samples. Plots of principal component 1 (PC1) vs principal component 2 (PC2) for gene expression variation in **A)** brain and **B)** blood samples, separated by age and pregnancy state of mice, respectively. The percentage of variance explained by each PC is indicated in the axis labels.

**Supplementary Figure 4:**
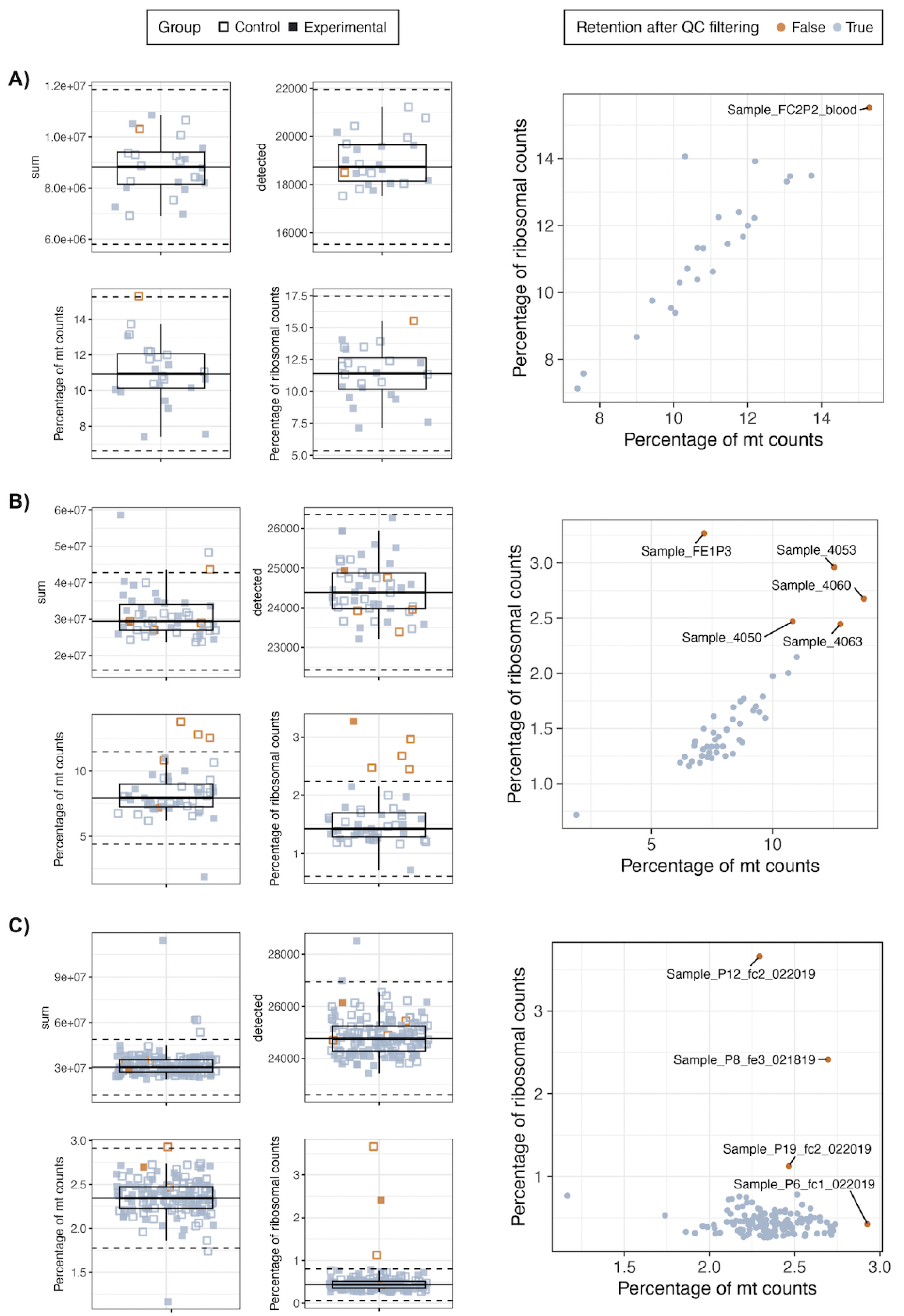
Sample filtering by QC metrics. Left box plots show the library size (top left), the total number of detected genes (top right), the percentage of the total sample counts that correspond to reads mapping to mitochondrial (bottom left) and ribosomal (bottom right) genes in **A)** adult blood, **B)** adult brain and **C)** pup brain samples. In orange, the samples that were removed after sample filtering based on such metrics; in blue the ones that passed the filtering step. `Group` separates samples in smoking/nicotine-exposed and smoking/nicotine controls. Dotted lines are 3 median-absolute-deviations away from the median (solid line) and set the cutoff values to determine if samples were or not taken as outliers; lower outlier samples in library size or detected number of genes, and higher outlier samples in mitochondrial or ribosomal percentages were considered poor-quality and thus discarded. The scatter plots on the right show the same percentages of mitochondrial (mt) and ribosomal genes’ read counts in all **A)** adult blood, **B)** adult brain and **C)** pup brain samples, labeling the filtered low-quality ones (in orange).

**Supplementary Figure 5:**
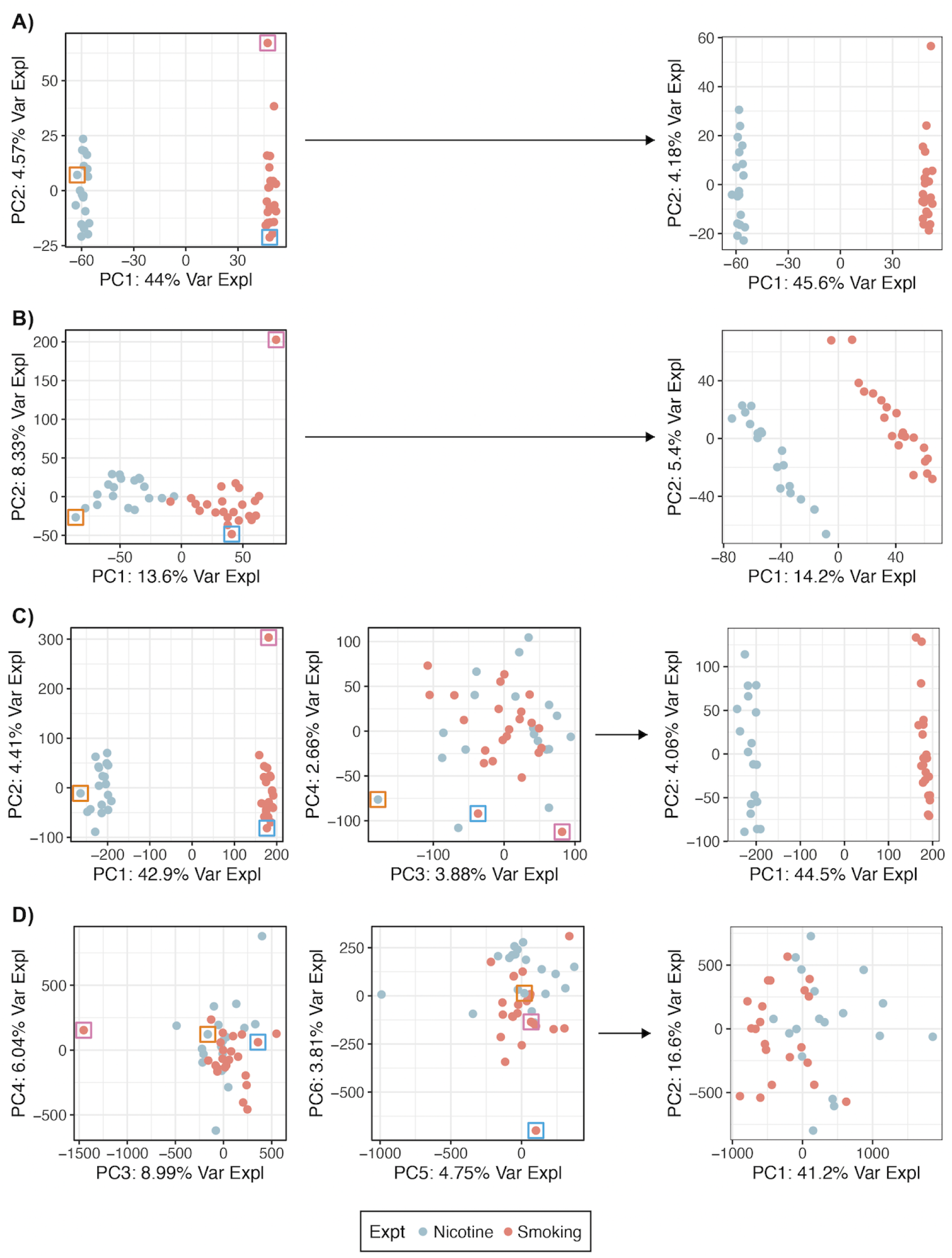
Manual sample filtering of adult brain samples. Plots of Principal Components (PCs) for **A)** gene, **B)** transcript, **C)** exon and **D)** exon-exon junction expression variation in brain samples from adult mice. In blue the samples from the nicotine experiment and in orange from the smoking experiment (both experiments include exposed and control samples). Left and middle plots contain all samples that passed QC-based sample filtering (see **Fig. S4**); right plots resulted from removing poor-quality samples that appeared very far from the rest in PC plots, boxed in different colors; dots boxed in the same color correspond to the same sample. ● Pink boxed sample: turned out to be the sample with the highest proportion of rRNA counts. ● Orange boxed sample: is the sample with the highest proportion of reads that mapped to the mitochondrial chromosome; it has the highest percentages of mitochondrial and ribosomal genes’ counts, the lowest proportion of reads assigned to genes and the minimum number of detected genes. ● Blue boxed sample: is the sample with the lowest decimal fraction of reads which successfully mapped to the reference genome and the smallest library size.

**Supplementary Figure 6:**
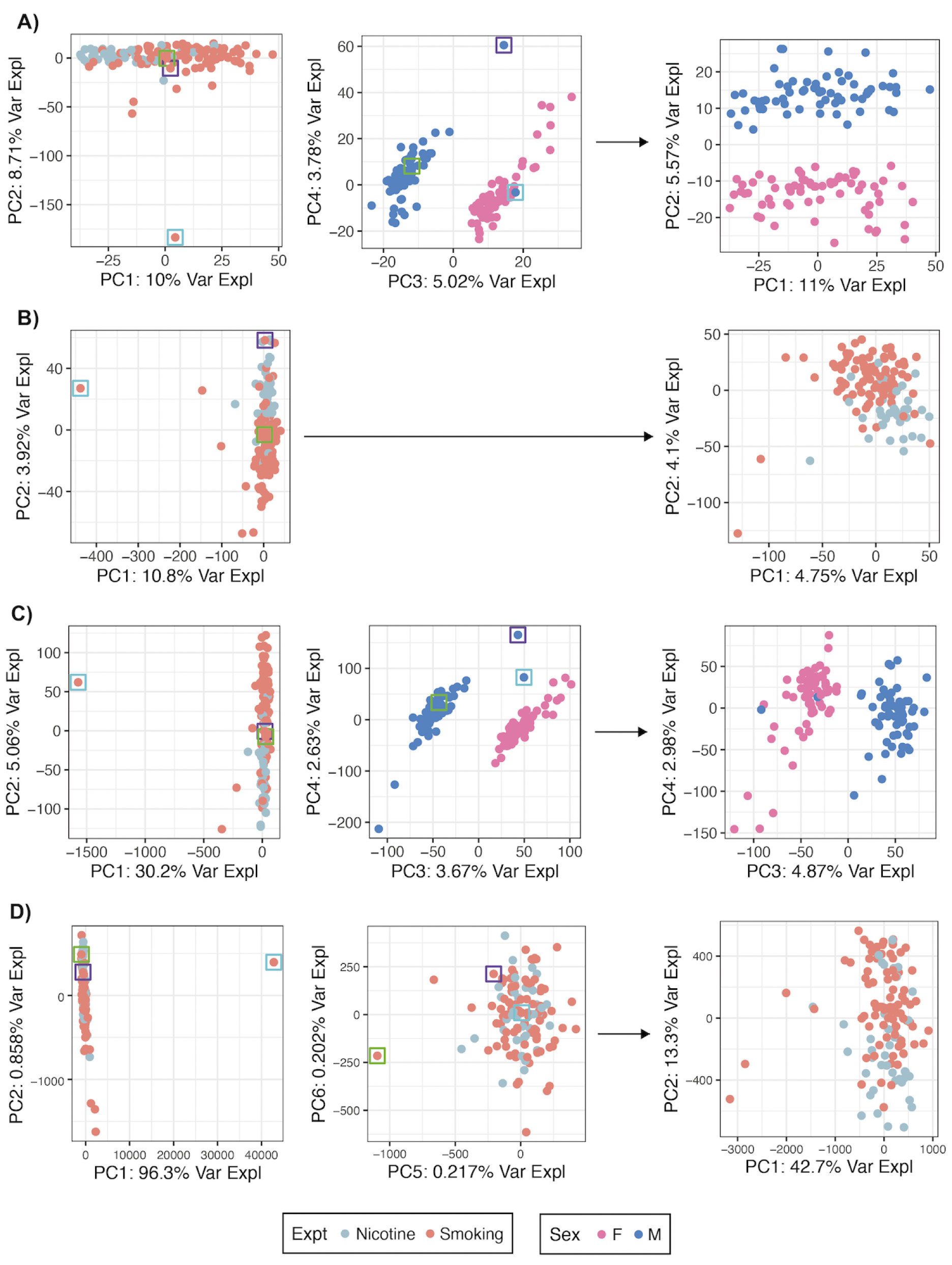
Manual sample filtering of pup brain samples. Plots of principal components for **A)** gene, **B)** transcript, **C)** exon and **D)** exon-exon junction expression variation in brain samples from pups. Left and middle plots contain all samples that passed QC-based filtering (see **Fig. S4**); right plots resulted from removing segregated samples in PC plots, those are boxed in the same color for the same sample. ‘Expt’ separates samples by experiment (PNE and MSDP, both including exposed and control samples) and ‘Sex’ into female (F) and male (M). ● Blue boxed sample: is the sample with the lowest proportion of reads assigned to genes. ● Purple boxed sample: is the sample with the highest overall difference between the expected and the actual External Control Consortium (ERCC) RNA concentrations. ● Green boxed sample: is the sample with the lowest decimal fraction of reads which successfully mapped to the reference genome and the smallest library size.

**Supplementary Figure 7:**
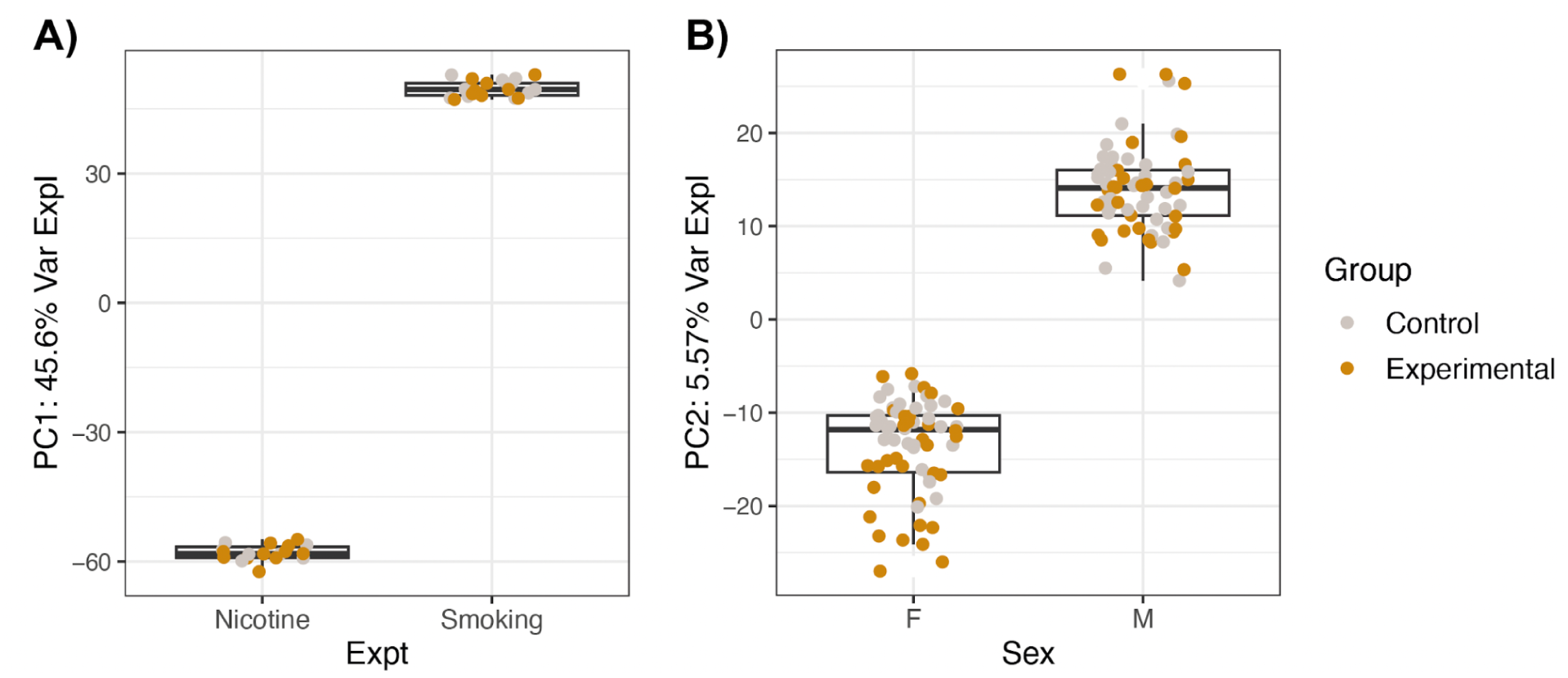
Explore gene expression variation in experimental and control brain samples. Box plots of principal components for gene expression variation in **A)** adult brain samples from mice of nicotine and smoking experiments and **B)** brain samples from female and male pups. Gray dots correspond to control samples and the brown ones to nicotine/smoking exposed samples.

**Supplementary Figure 8:**
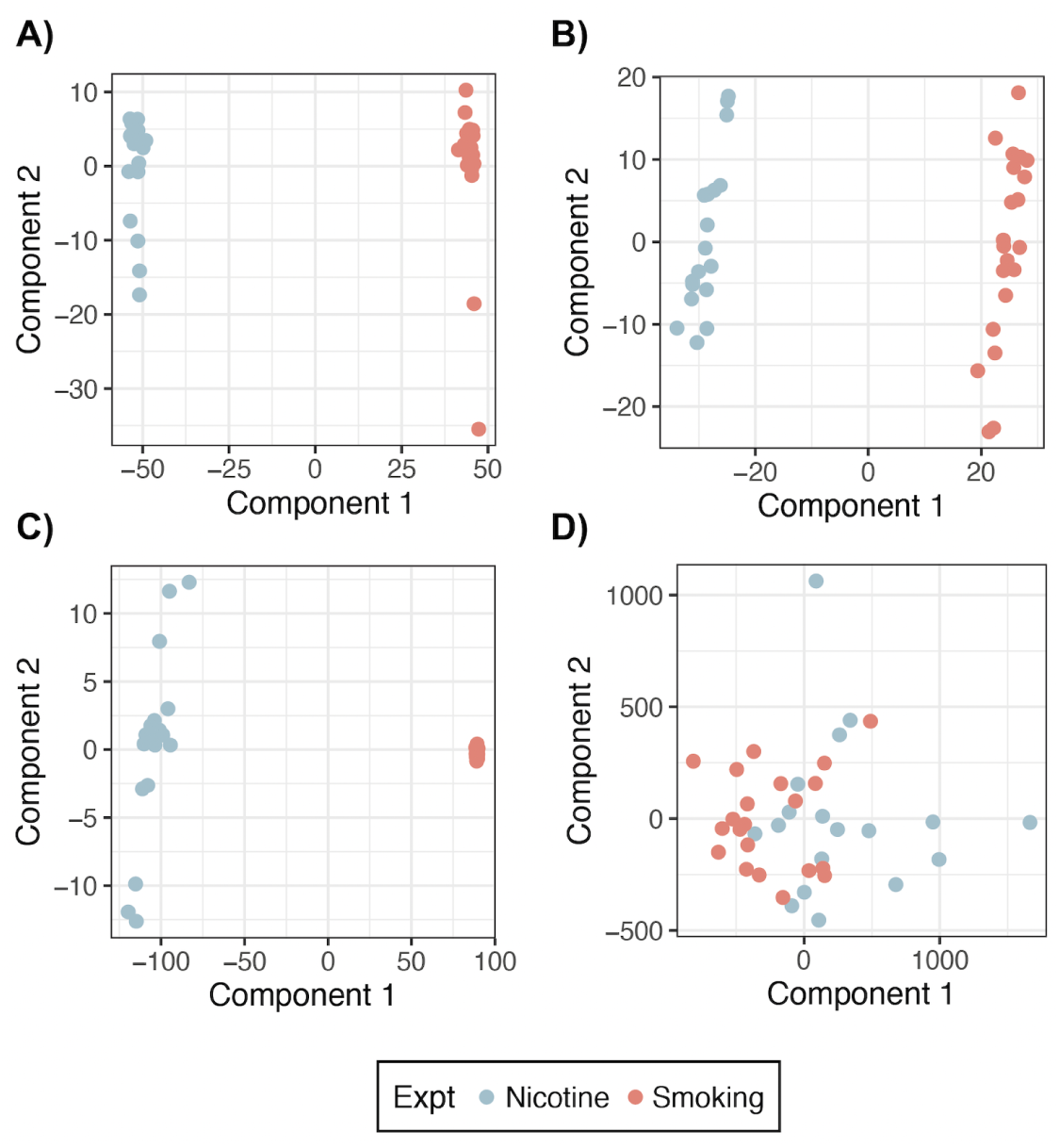
Multidimensional scaling analysis in filtered adult brain samples. Component 1 vs Component 2 for **A)** gene, **B)** transcript, **C)** exon and **D)** exon-exon junction expression variation in adult brain samples from the nicotine and smoking experiments, including exposed and control samples. This analysis was done with samples that passed QC and manual sample filtering only (see **Fig. S4** and **Fig. S5**).

**Supplementary Figure 9:**
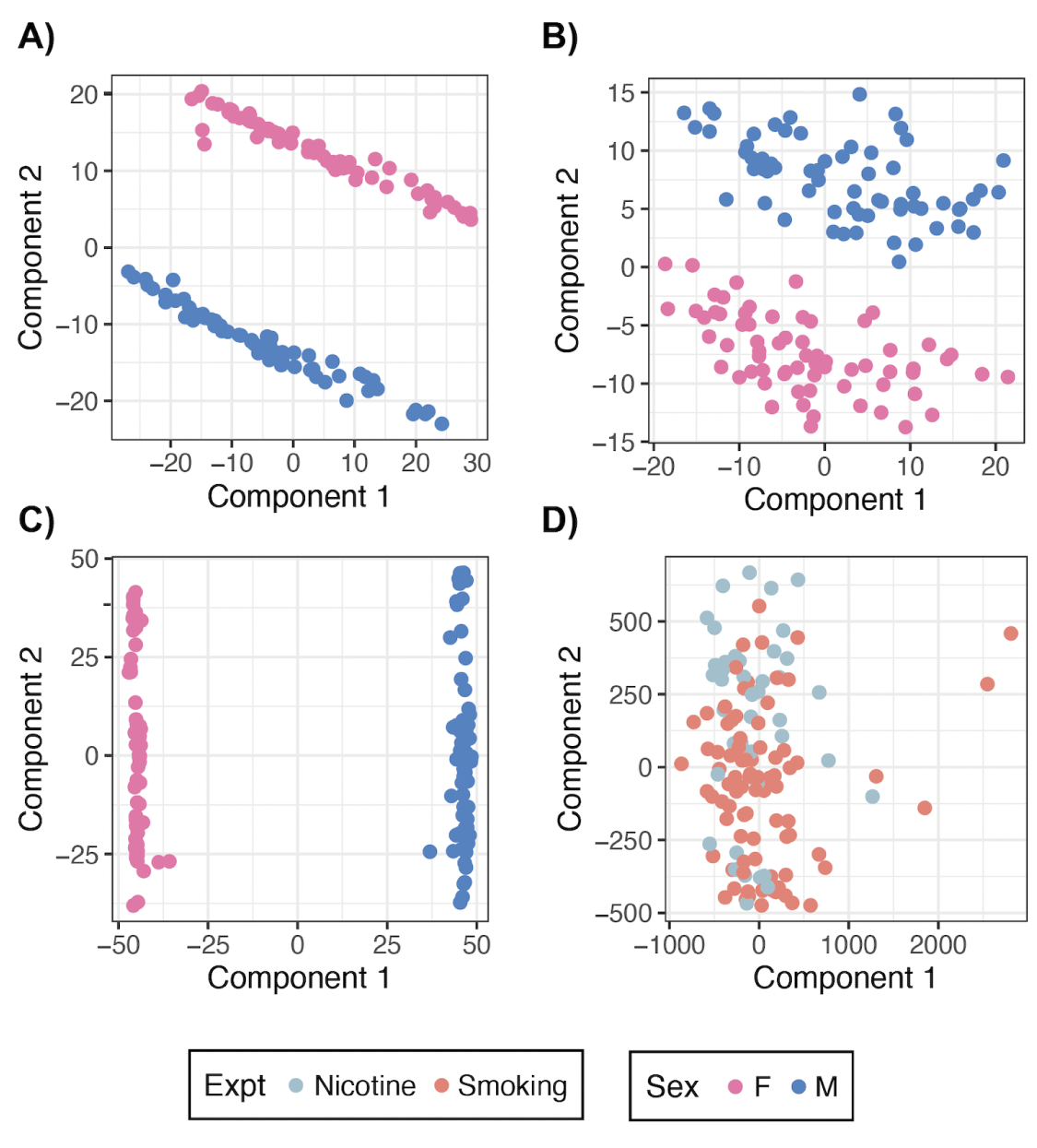
Multidimensional scaling analysis in filtered pup brain samples. Component 1 vs Component 2 for **A)** gene, **B)** transcript, **C)** exon and **D)** exon-exon junction expression variation in pup brain samples that passed QC and manual sample filtering (see **Fig. S4** and **Fig. S6**). In **A)**-**C)** samples are separated by sex: females (F) and males (M); in **D)** by experiment, including exposed and control samples.

**Supplementary Figure 10:**
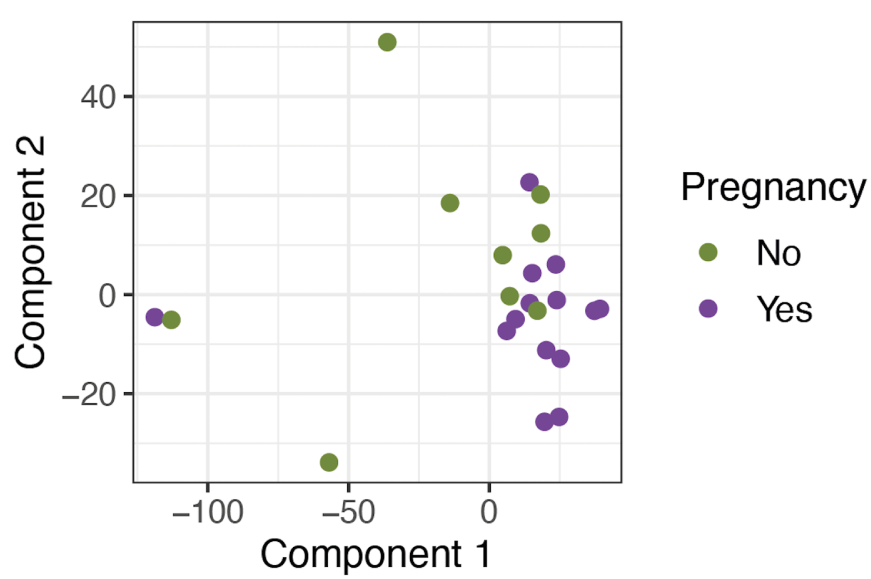
Multidimensional scaling analysis in filtered blood samples. Component 1 vs Component 2 for gene expression variation in blood samples from pregnant and non-pregnant mice; these correspond to samples that passed QC sample filtering (see **Fig. S4**).

**Supplementary Figure 11:**
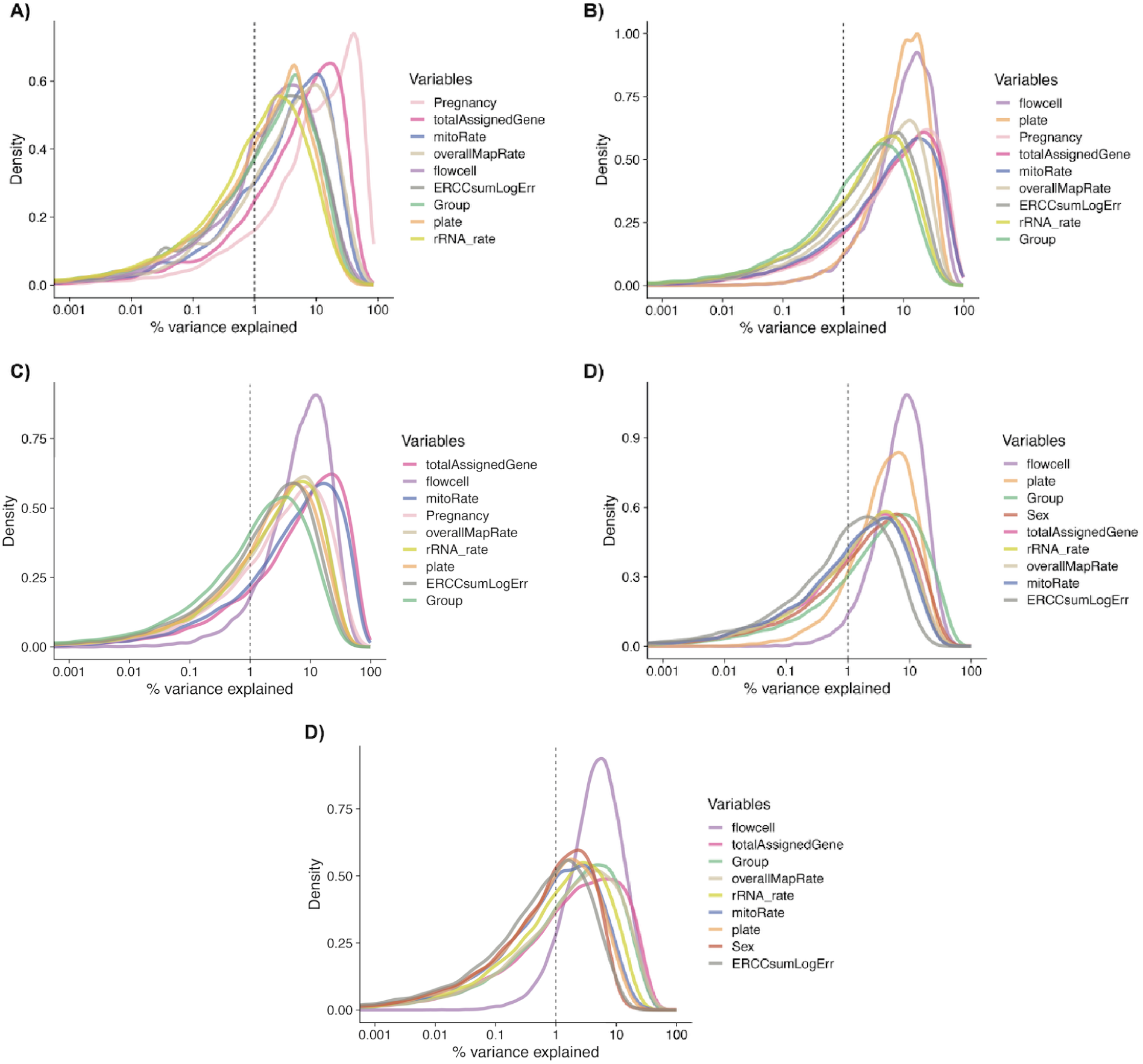
Analysis of variance in gene expression explained by the explanatory sample variables. Density plots for the percentage of variance in the expression of each gene that is explained by each sample-level variable in **A)** blood samples, **B)** adult brain samples from the nicotine or **C)** smoking experiment and **D)** pup brain samples from the nicotine or **E)** smoking experiment. See **Table S18** for the description of the covariates.

**Supplementary Figure 12:**
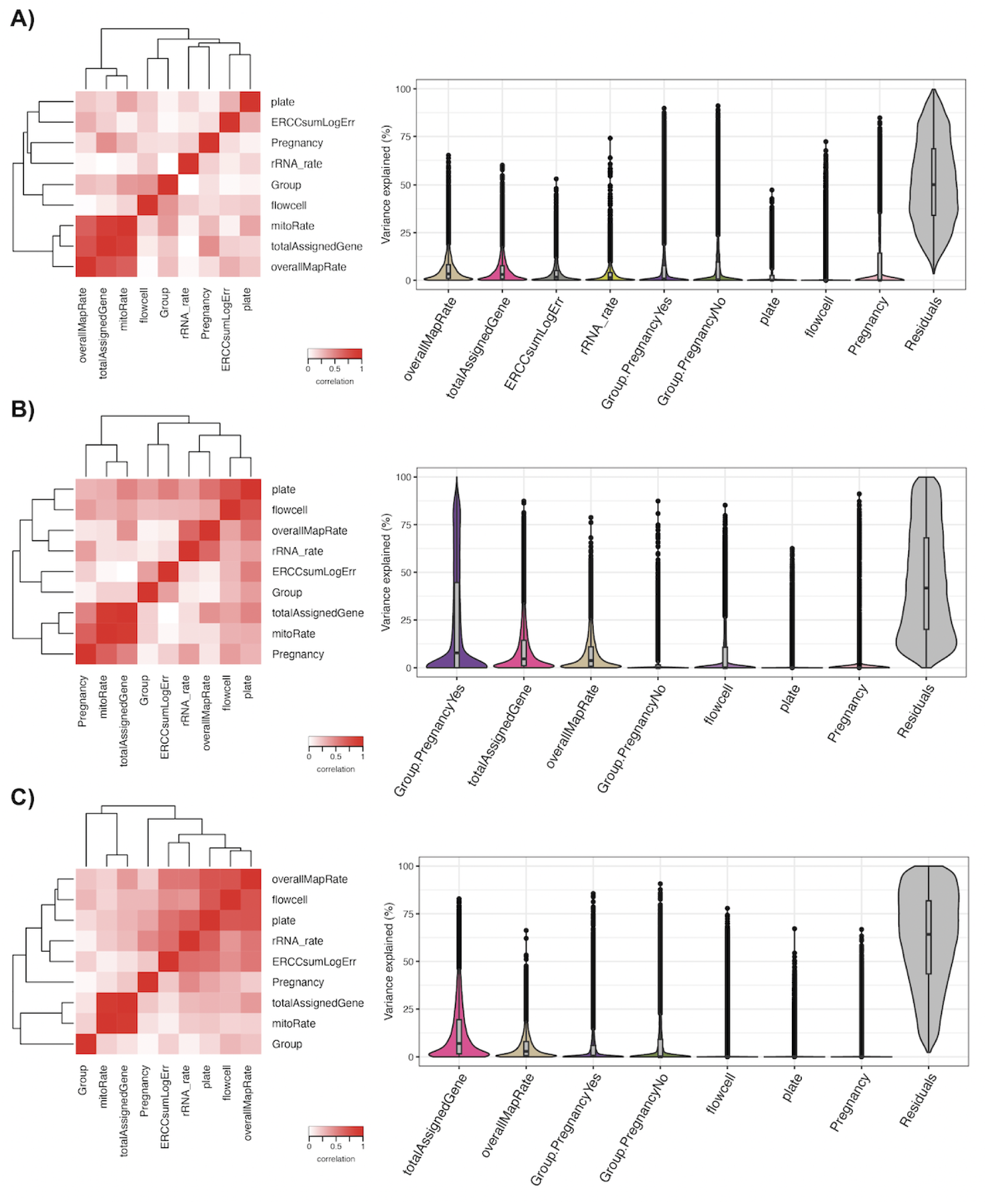
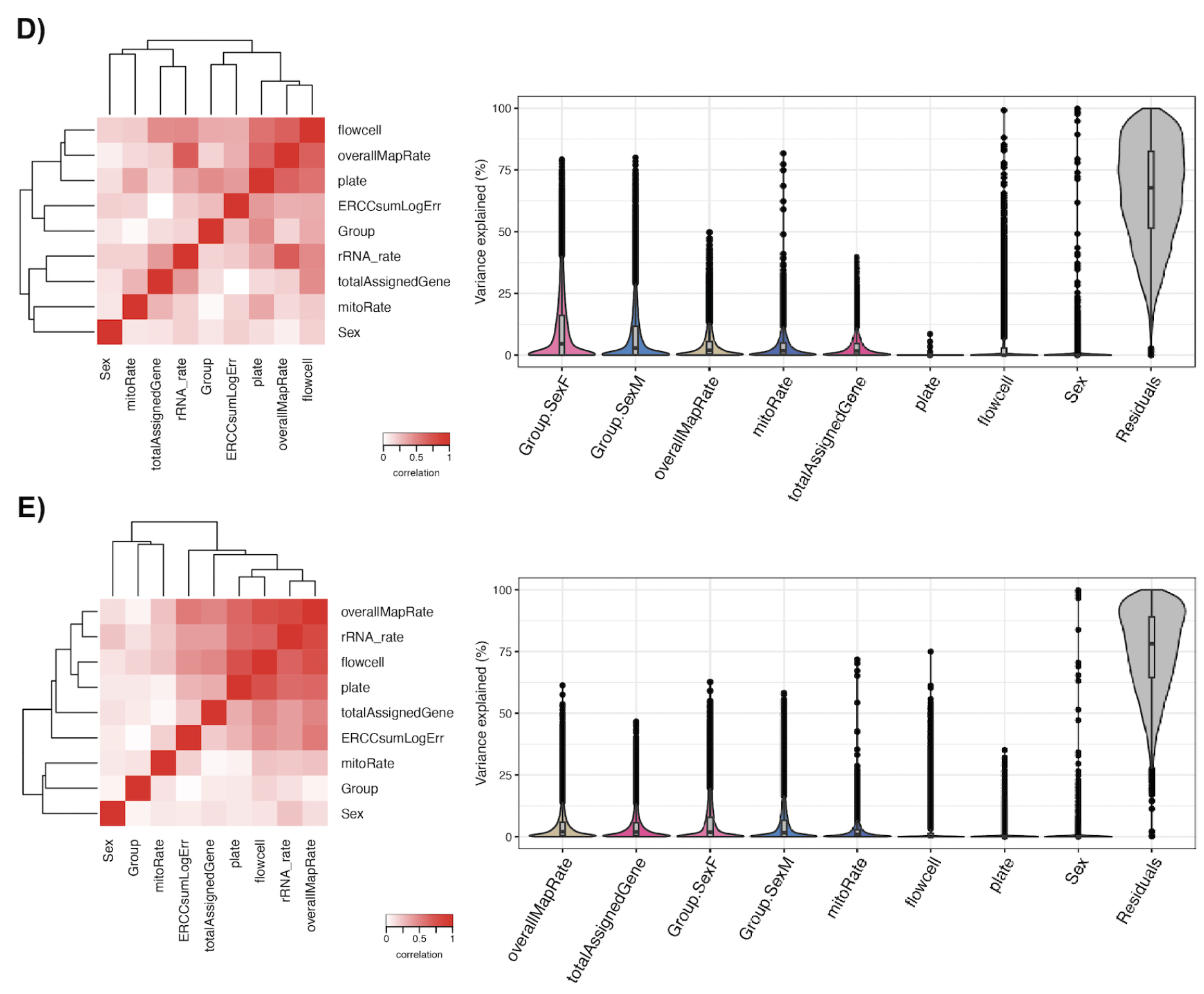
Variance partition analysis. Left heat maps show the correlation (from 0 to 1) between each pair of sample variables, obtained through a Canonical Correlation Analysis (CCA). Violin plots show for each gene the percentages of variance in their expression levels that are explained by each variable in **A)** blood samples of the smoking experiment, **B)** adult brain samples of the nicotine experiment, **C)** adult brain samples of the smoking experiment, **D)** pup brain samples of the nicotine experiment, and **E)** pup brain samples of the smoking experiment. Variables are ordered by decreasing mean fraction of variance explained (FVE). Variables in the heat maps not present in the corresponding violin plots were highly correlated with any other variable with a higher median FVE and thus were not included in the models for DGE. `ERCCsumLogErr` and `rRNA_rate` were not considered in the brain samples because their scales differ considerably and therefore were not suitable for variance partition. Labels `Group:Pregnancy` in **A)**-**C)** and `Group:Sex` in **D)** and **E)** refer to the interaction of nicotine/smoking exposure (`Group`) with pregnancy and sex, respectively. See **Table S18** for the description of these sample variables. Residuals correspond to those fractions of gene expression variation that could not be attributed to any of the sample-level variables.

**Supplementary Figure 13:**
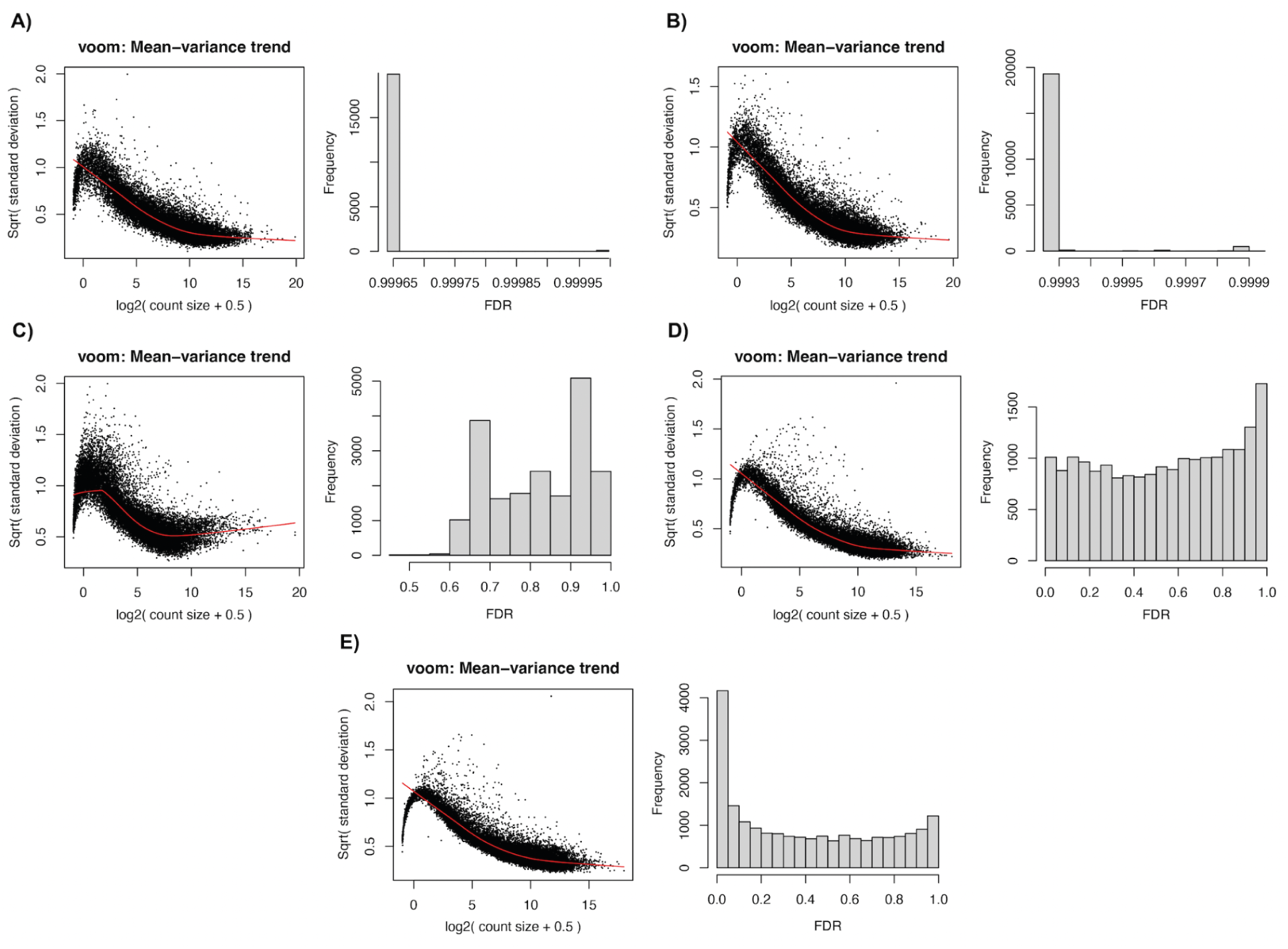
**Results of differential gene expression analyses**. Left plots show for each gene their mean expression (in log2-counts) and the square-root of their residual standard deviations in **A)** adult brain samples of the nicotine experiment, **B)** adult brain samples of the smoking experiment, **C)** adult blood samples of the smoking experiment, **D)** pup brain samples of the nicotine experiment, and **E)** pup brain samples of the smoking experiment. Red line corresponds to the global mean-variance trend. Histograms present gene-wise FDR-adjusted *p*-values for differential expression in the same sample groups.

**Supplementary Figure 14:**
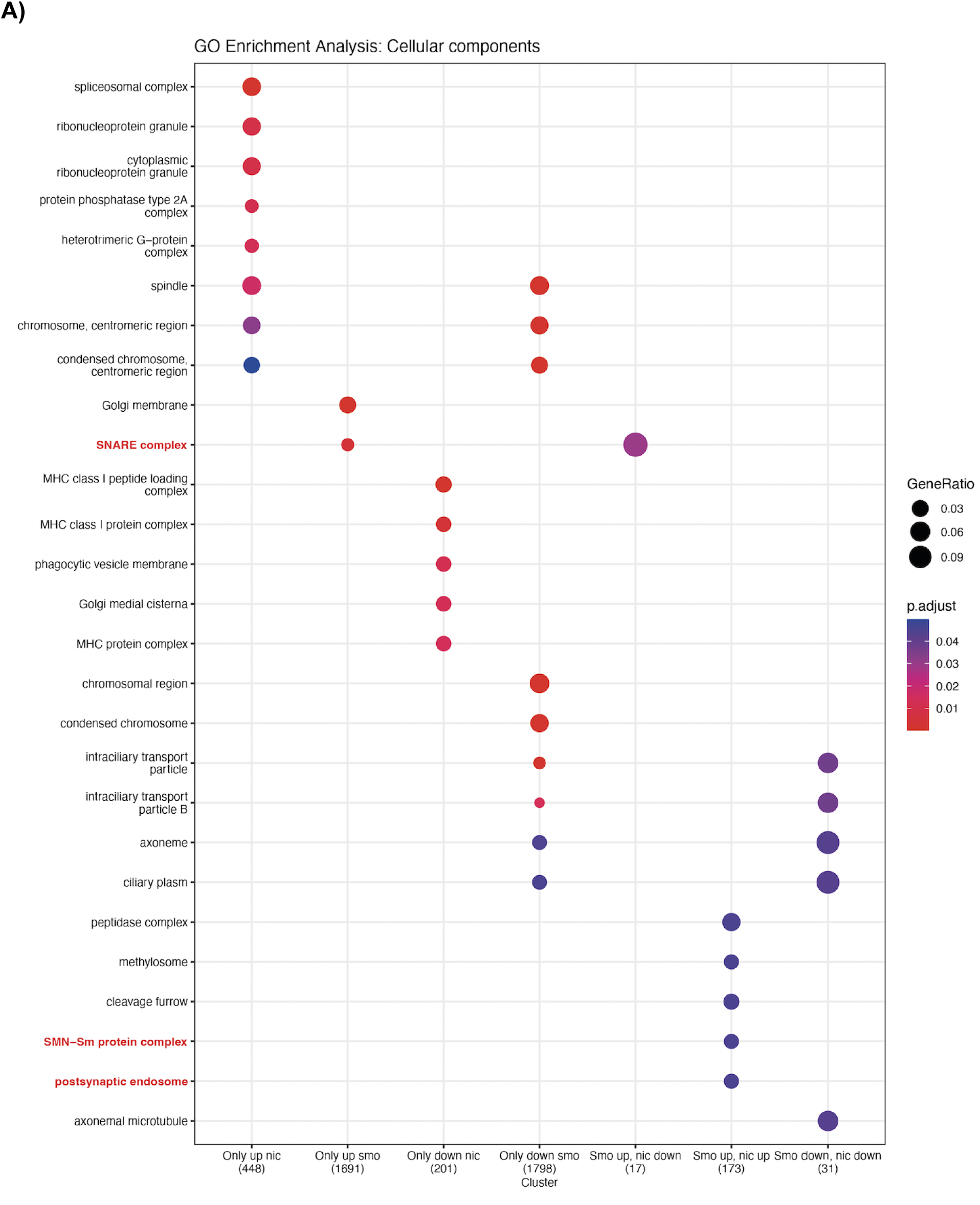
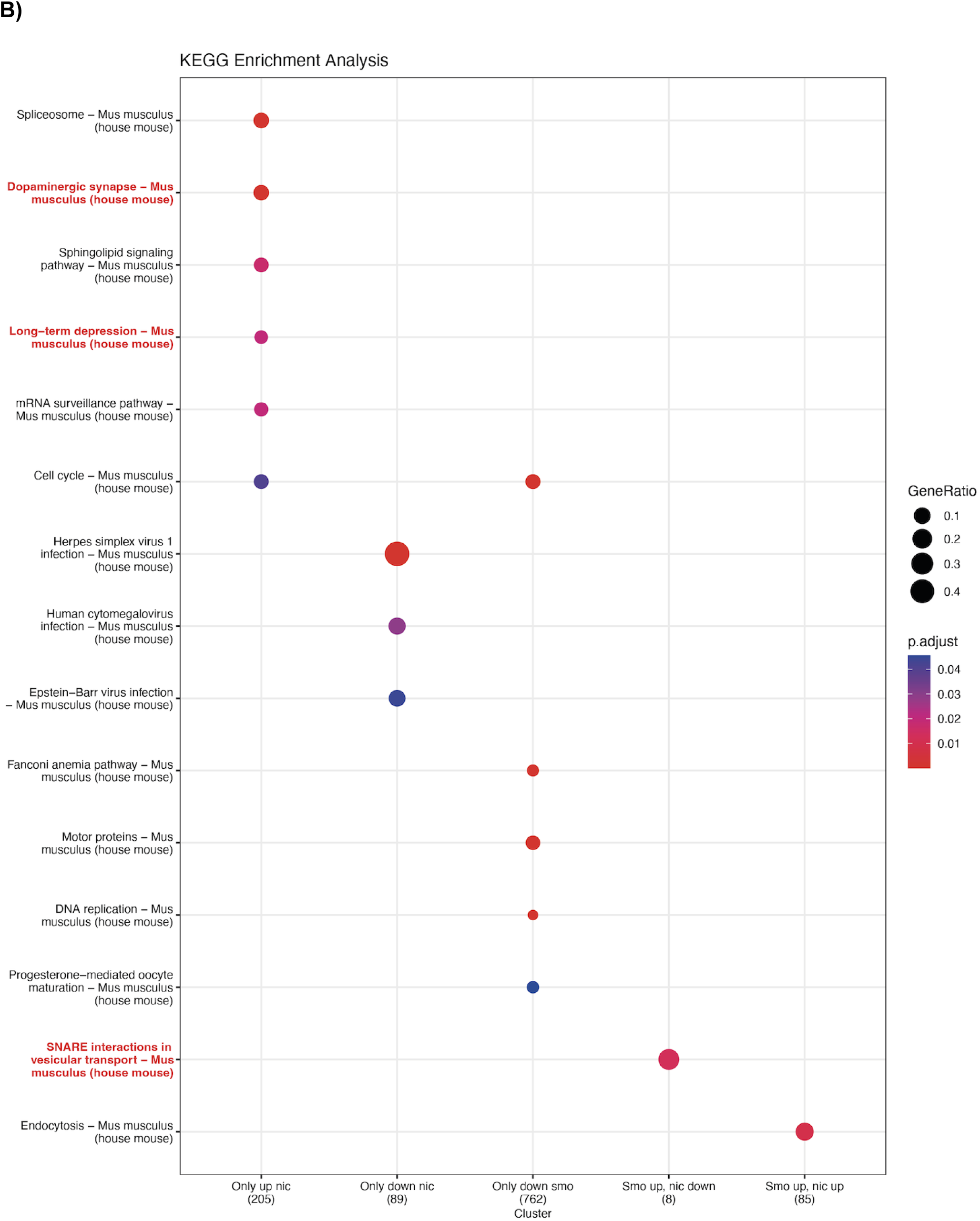
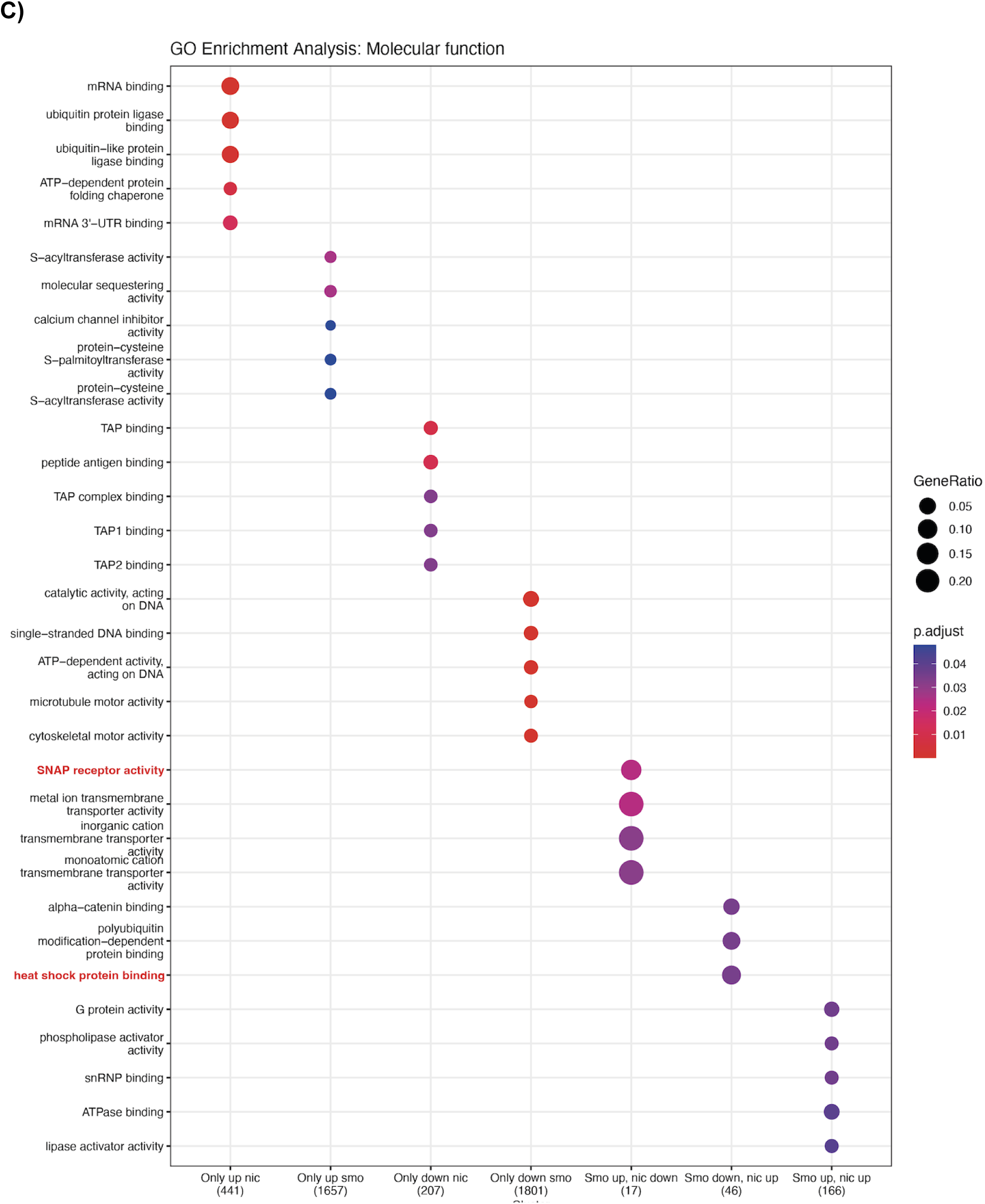
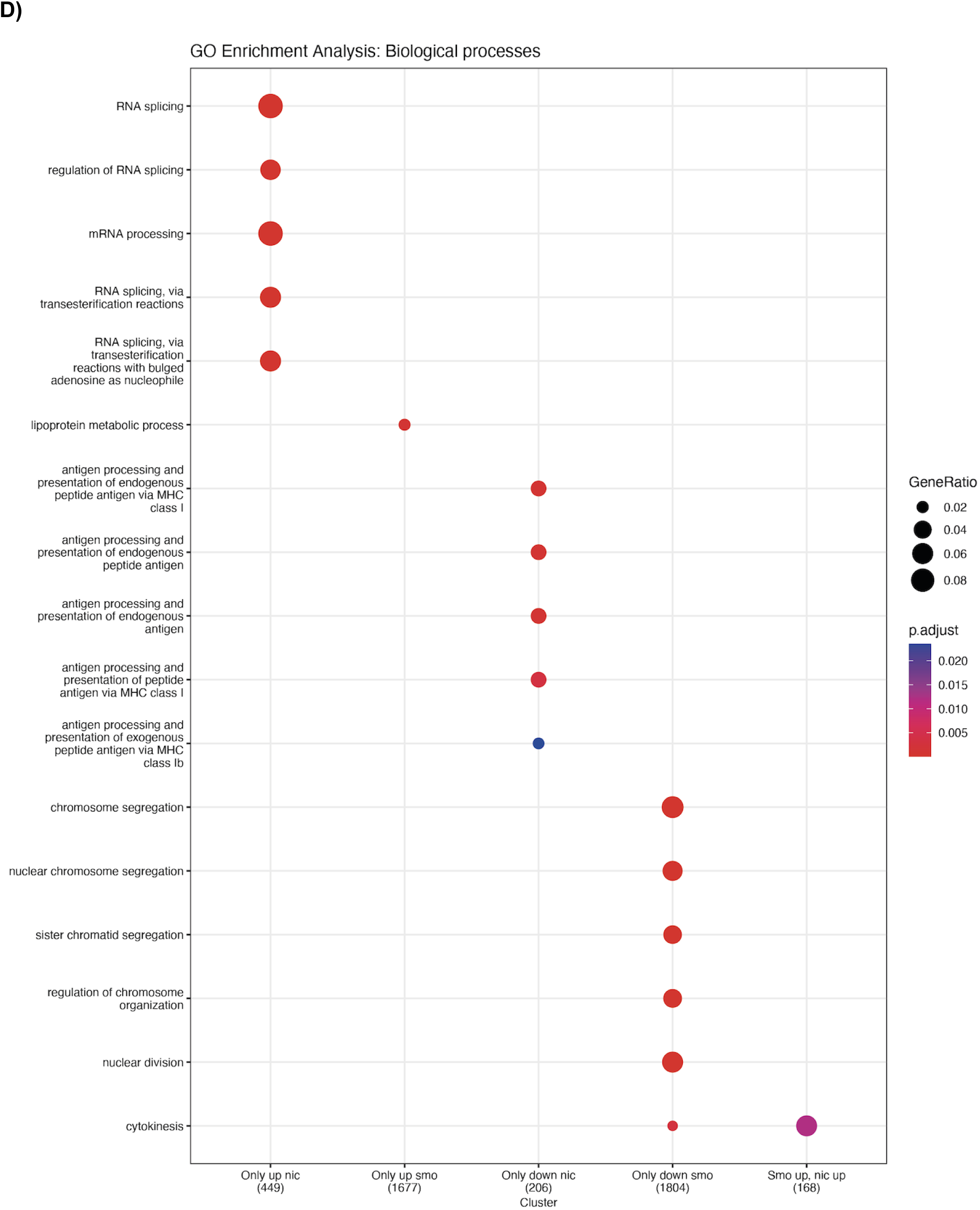
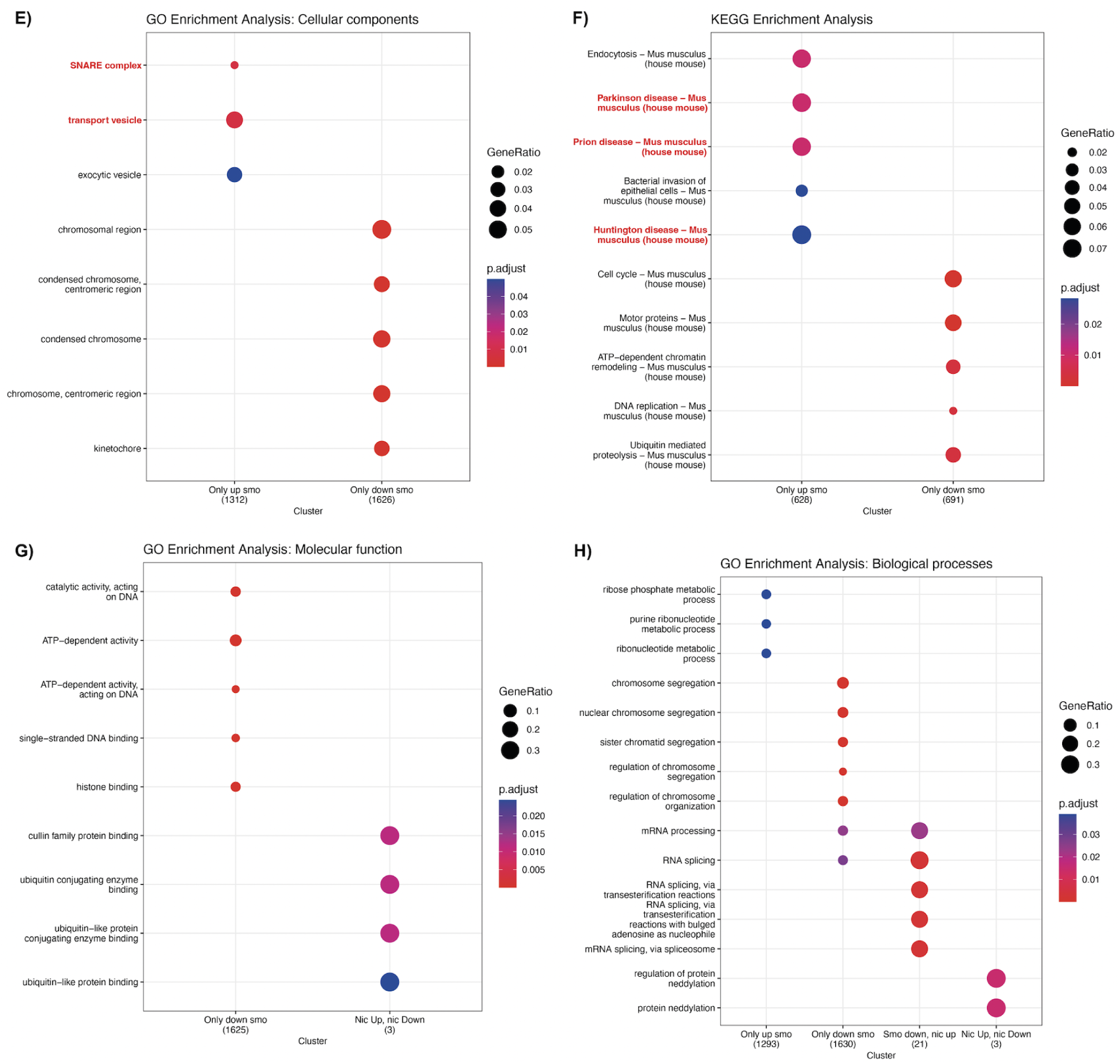
**Functional enrichment analysis for DEGs and DE transcripts’ genes in pup brain**. Cellular components (CC), pathways, molecular functions (MF) and biological processes (BP) significantly overrepresented (FDR-adjusted p-value<0.05) in the clusters of (**A-D**) DEGs and (**E-H**) genes with DE transcripts, indicated in the x-axis. Clusters without significant results are excluded; `up` and `down` labels stand for upregulated and downregulated, respectively, and `only` refers to genes that were significant (**A-D**) or had significant transcripts (**E-H**) in either the smoking (`smo`) or nicotine (`nic`) experiment but not in the other. Note that cluster numbers (in parentheses) correspond to the number of genes in the specified cluster that are annotated in at least one GO/KEGG term. Gene ratio is the number of genes in each cluster annotated in a term over the total in the respective cluster. Only the top 5 most significant enriched terms are reported per cluster, unless they share additional significant terms with other clusters. Terms of interest appear in red (see genes involved in each in **Fig. S15**). Related to Fig. 2.

**Supplementary Figure 15:**
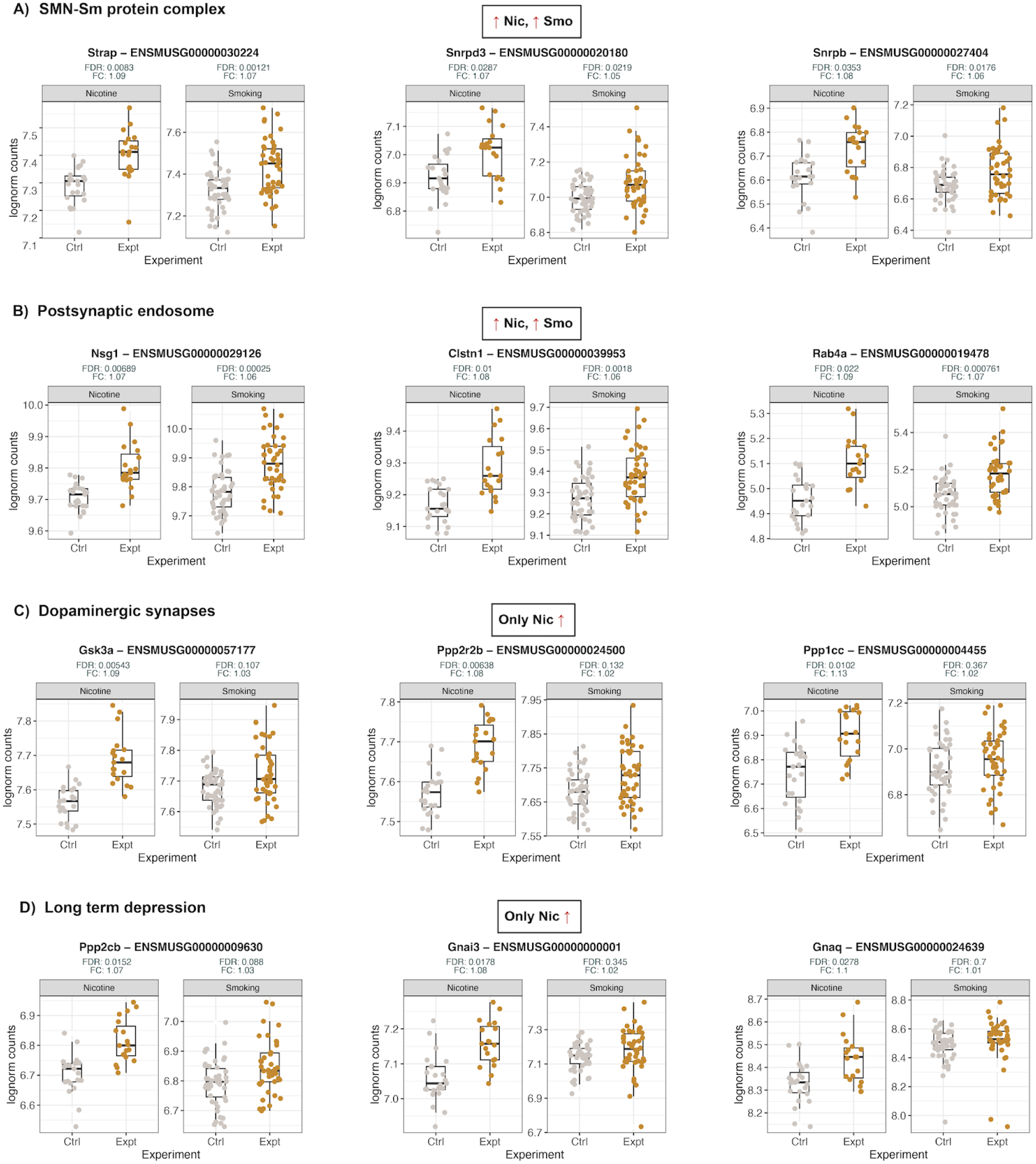
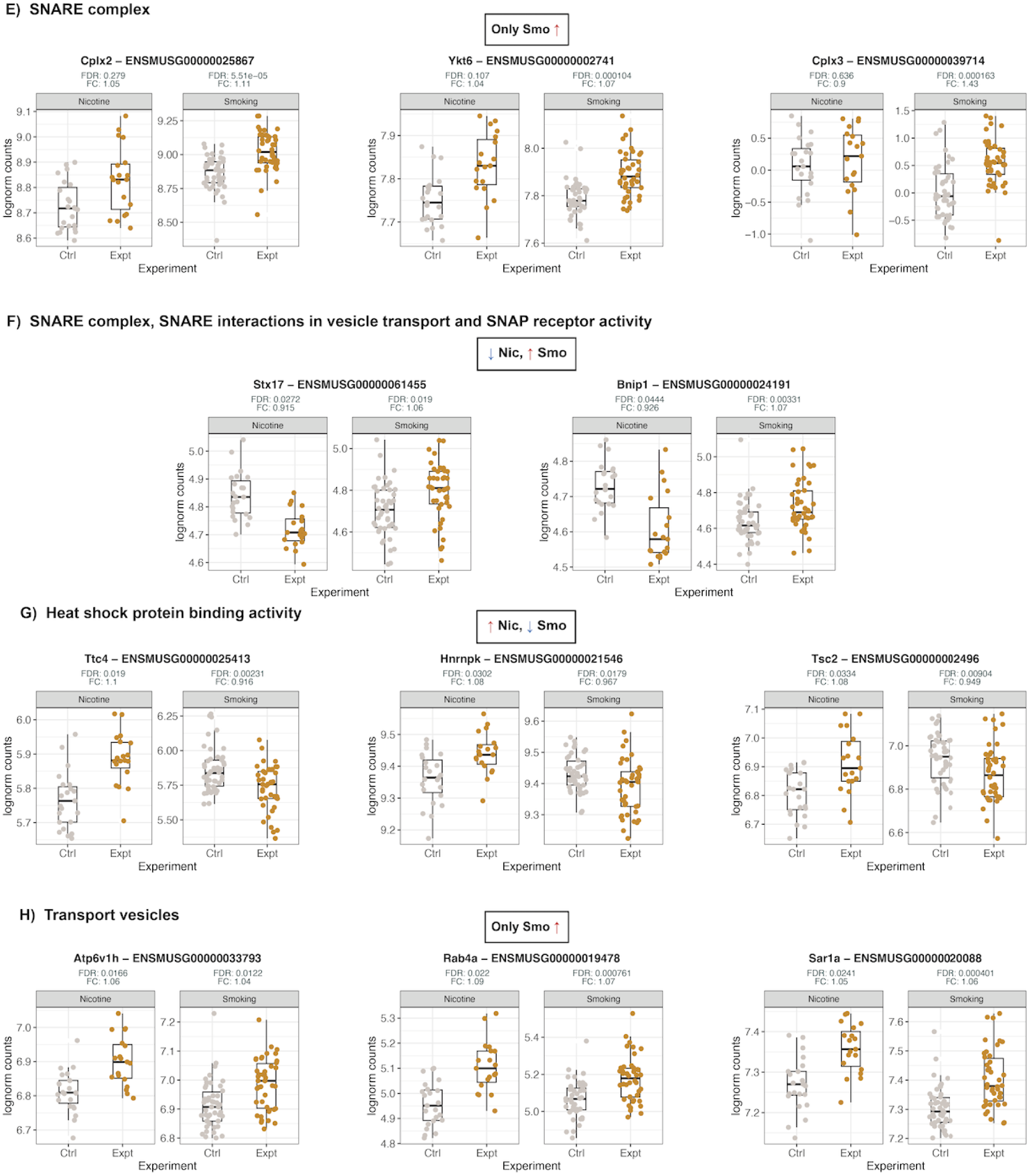
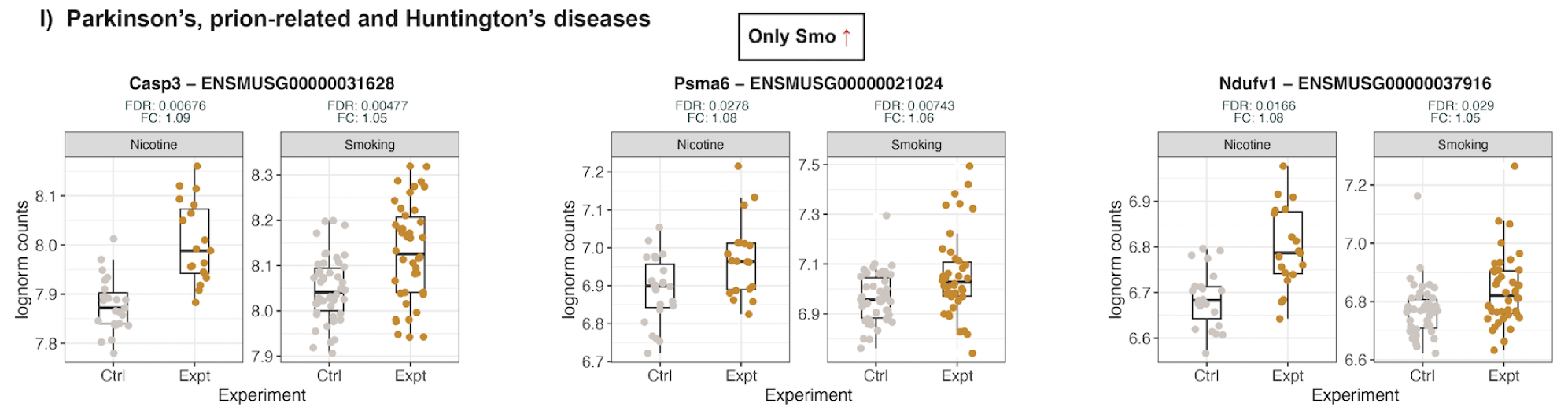
**Expression of genes implicated in interest brain-related processes in pup brain**. Gene lognorm-counts of **A)** the top 3 most significant DEGs upregulated in nicotine and smoking-exposed samples that act in SMN-Sm protein complexes and **B)** postsynaptic endosomes. **C)** The top 3 most significant DEGs upregulated in nicotine samples only that are involved in dopaminergic synapses and **D)** in long-term depression. **E)** The top 3 most significant DEGs whose products work in the SNARE complex and are upregulated in smoking samples only (their DE transcripts too) or **F)** are up in smoking and down in the nicotine experiment and also have SNAP receptor activity and are implicated in SNARE interactions in vesicle transport. **G)** The top 3 DEGs upregulated for nicotine and downregulated for smoking whose gene products have heat shock protein binding activity. **H)** The top 3 most significant genes with DE transcripts that were upregulated only by cigarette smoke and act in transport vesicles and that are involved in **I)** Parkinson’s, prion-related, and Huntington’s diseases. Related to Fig. 2 and **Fig. S14**. FDR: false discovery rate; FC: fold-change; Ctrl: control samples; Expt: experimental (exposed) samples.

**Supplementary Figure 16:**
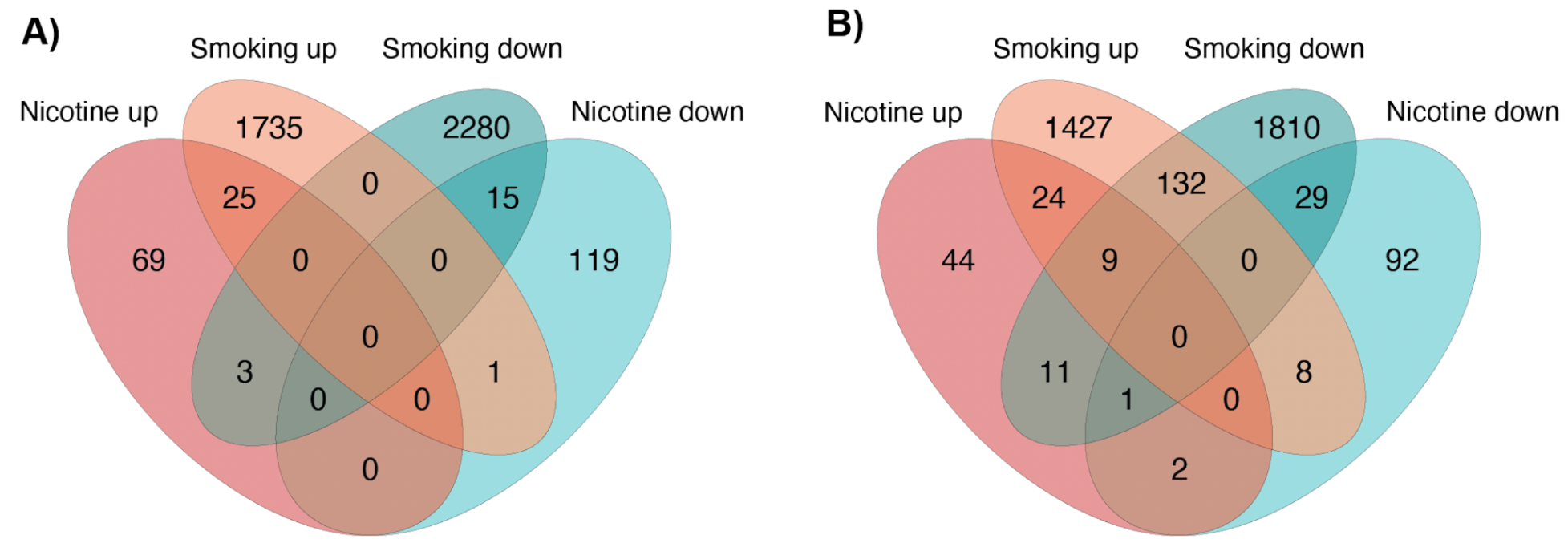
Results of the DTE analysis in pup brain. Number of **A)** DE transcripts and **B)** genes with DE transcripts, up- and down-regulated in the nicotine and smoking experiments.

**Supplementary Figure 17:**
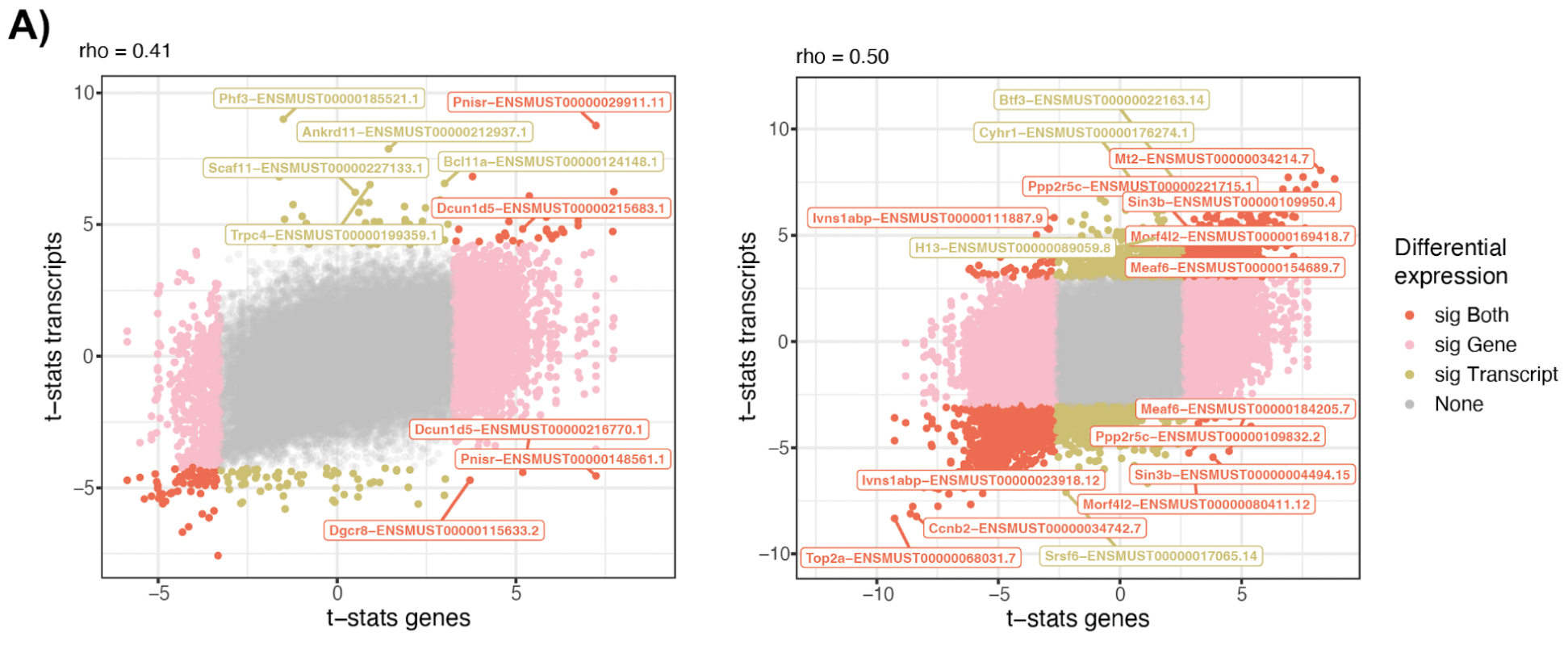
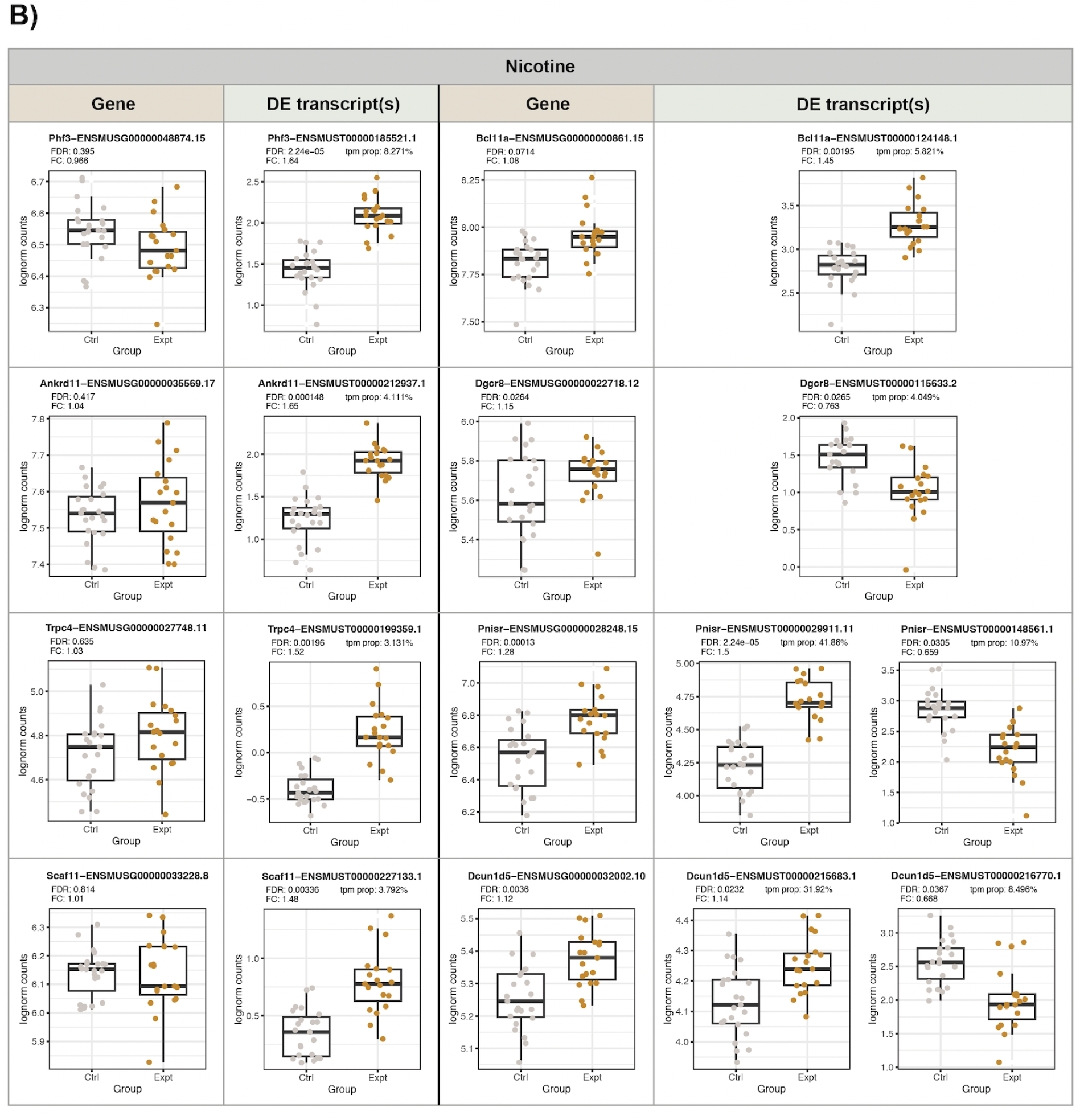
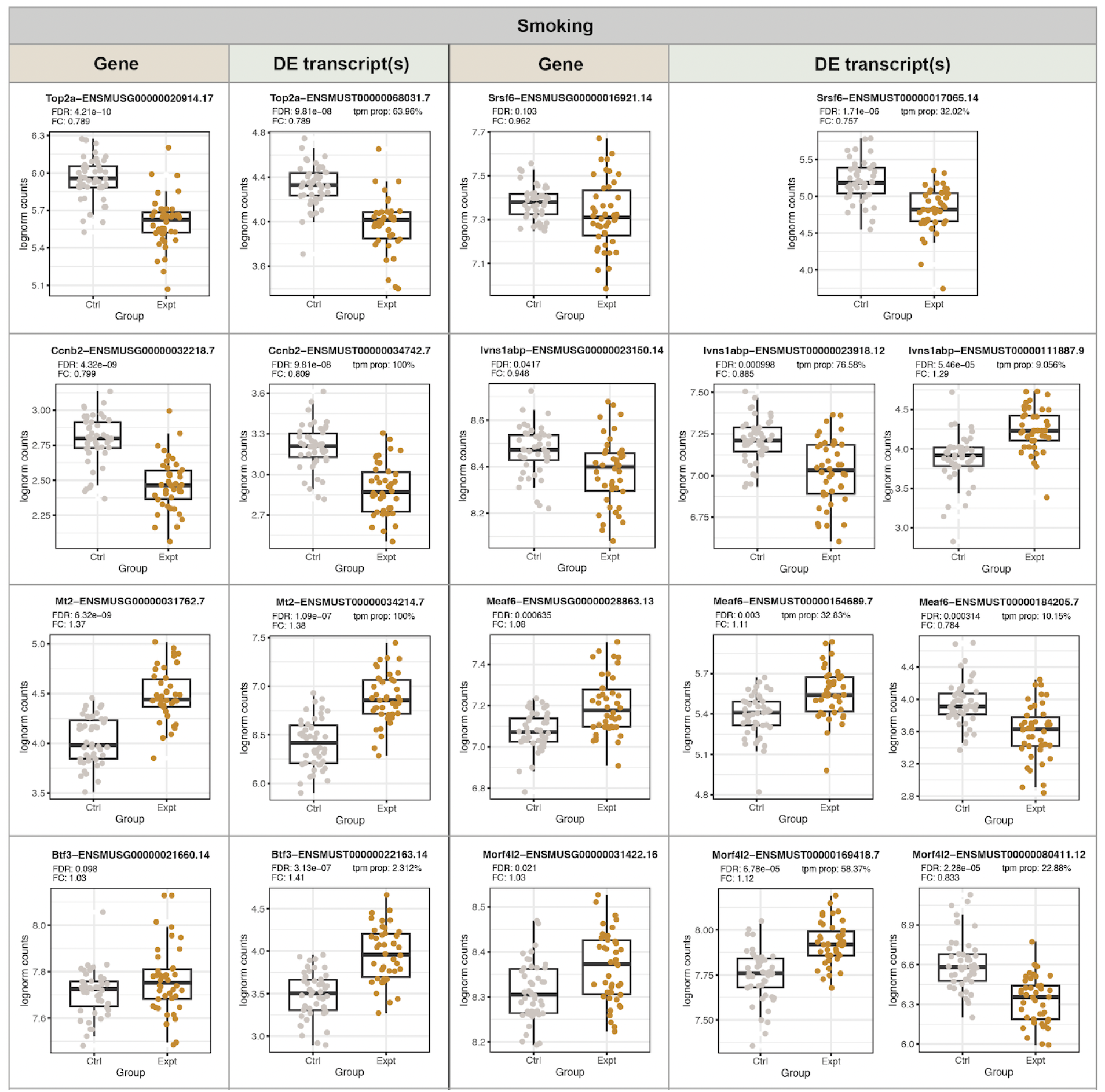
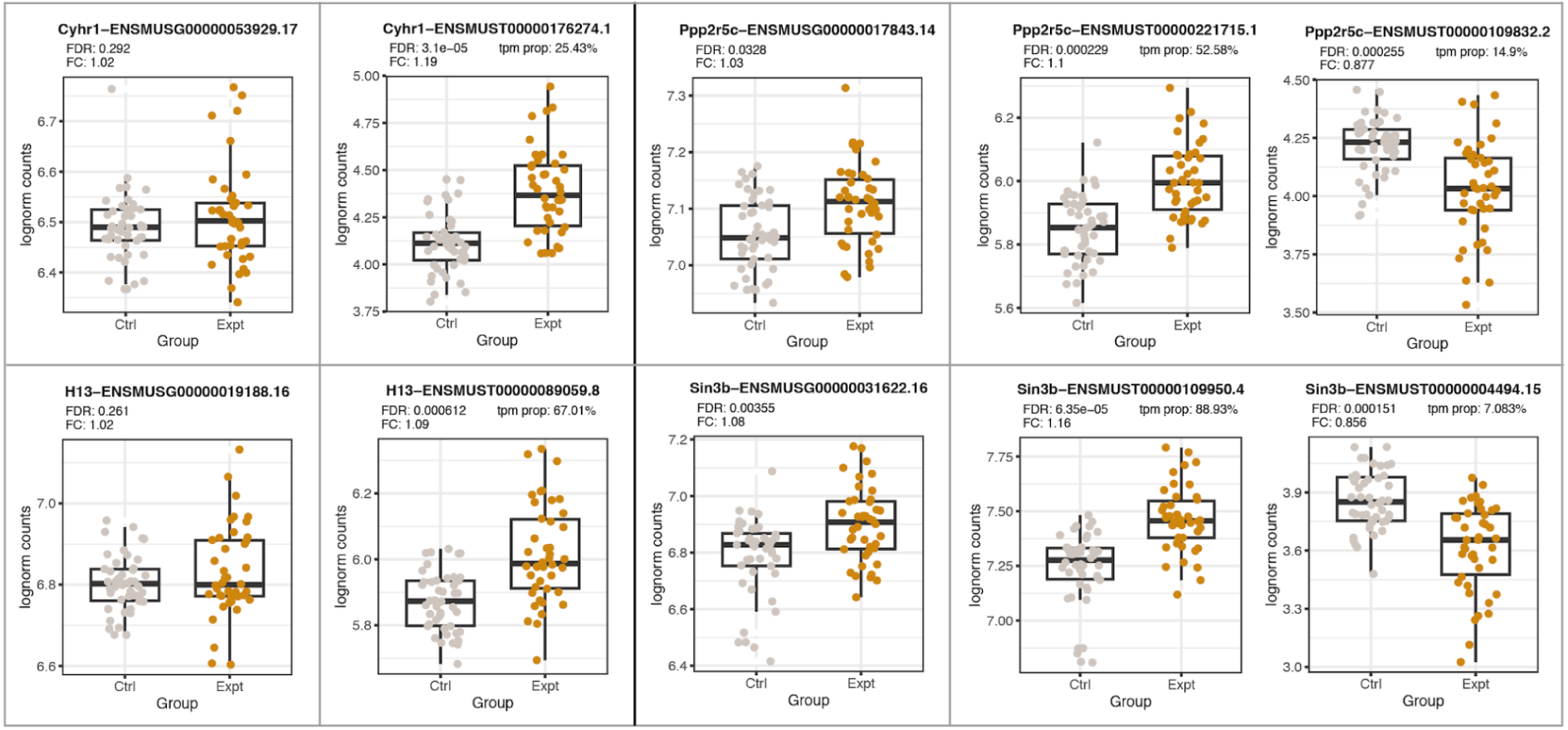
Expression of DE transcripts and their genes. **A)** Plots showing the differential expression results as in Fig. 3A for prenatal nicotine vs vehicle exposure (left), and prenatal smoking exposure vs control (right). The significant DE transcripts of interest are labeled with their corresponding gene symbol and transcript Ensembl ID. **B)** Box plots show the expression of the labeled DE transcripts (in log-tpm) and their corresponding genes (in log-cpm) for the nicotine and smoking exposure. FDR: false discovery rate; FC: fold-change; tpm prop: the proportion of the total TPM of a gene that corresponds to the transcript. Total gene TPM was obtained adding TPM of all transcripts of the gene across all samples.

**Supplementary Figure 18:**
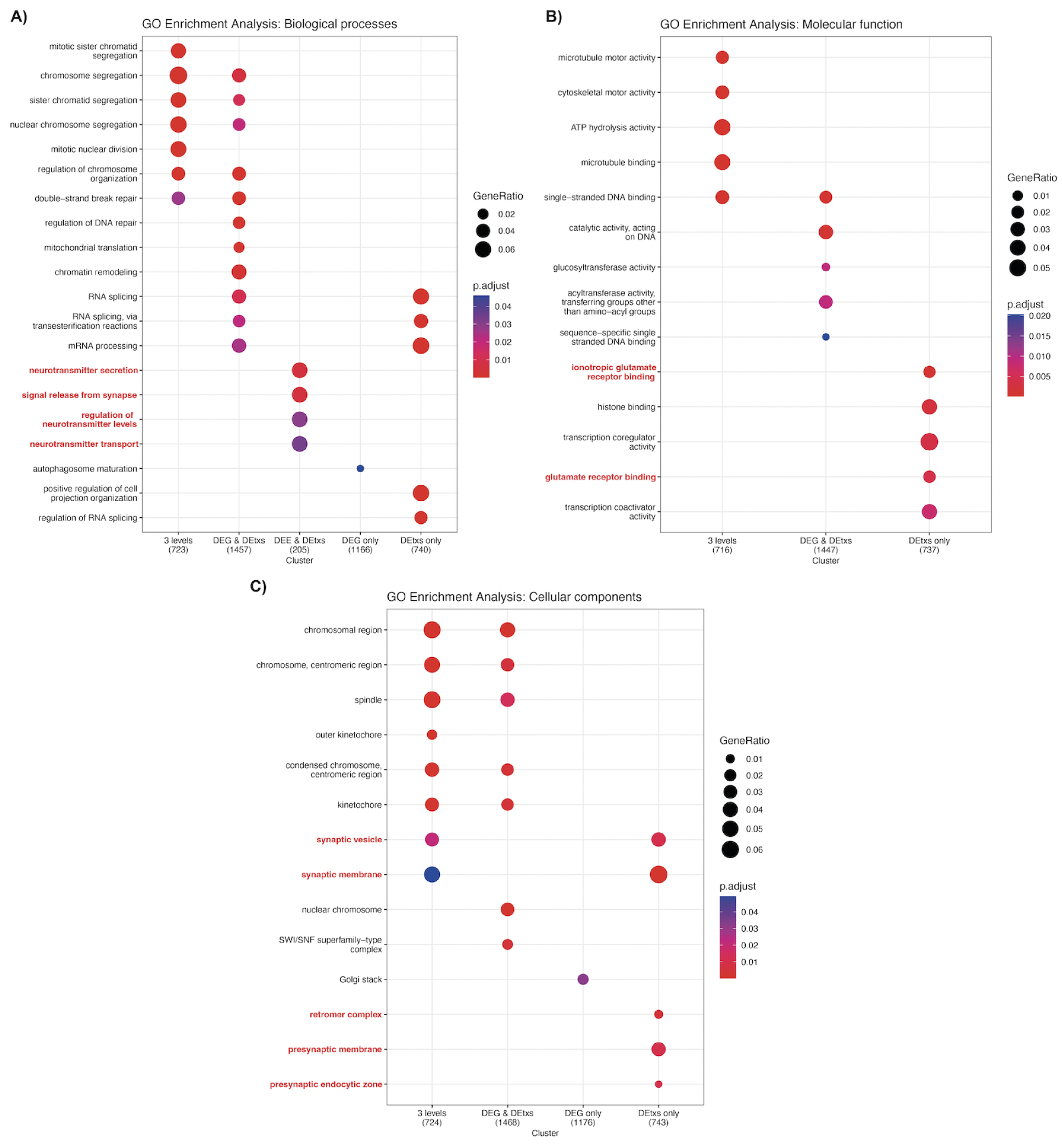
**Functional enrichment analysis for genes with DE features in pup brain**. **A)** Biological processes, **B)** molecular functions, and **C)** cellular components significantly enriched (adjusted p-value<0.05) in clusters of DEGs with DE transcripts and DE exons (`3 levels`), DEGs with DE transcripts (`DEG & DEtxs`), DEGs with DE exons (`DEG & DEE`), non-DEGs with DE transcripts and DE exons (`DEE & DEtxs`), DEGs only (`DEG only`), and non-DEGs with DE transcripts only (`DEtxs only`) or DE exons only (`DEE only`) for the smoking experiment. The terms of interest appear in red. See **Fig. S14** caption for more details of these plots.

**Supplementary Figure 19:**
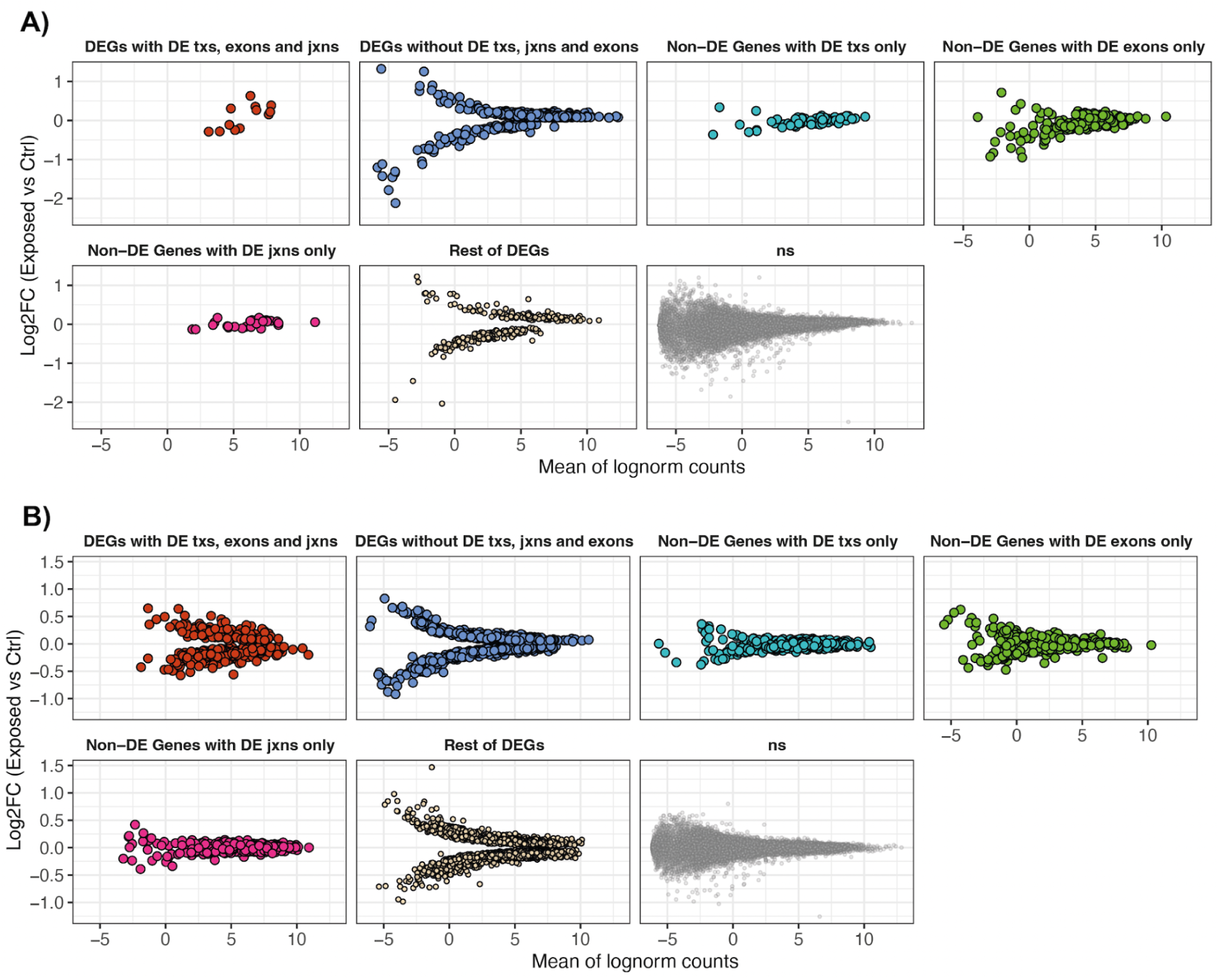
MA plots of genes DE at different expression levels. Mean log-cpm and logFC of DEGs and non-DEGs with and without DE transcripts (txs), exons, and exon-exon junctions (jxns) for the **A)** nicotine and **B)** smoking exposure. Rest of DEGs are DEGs with two other DE features (txs and exons, txs and jxns, or exons and jxns).

**Supplementary Figure 20:**
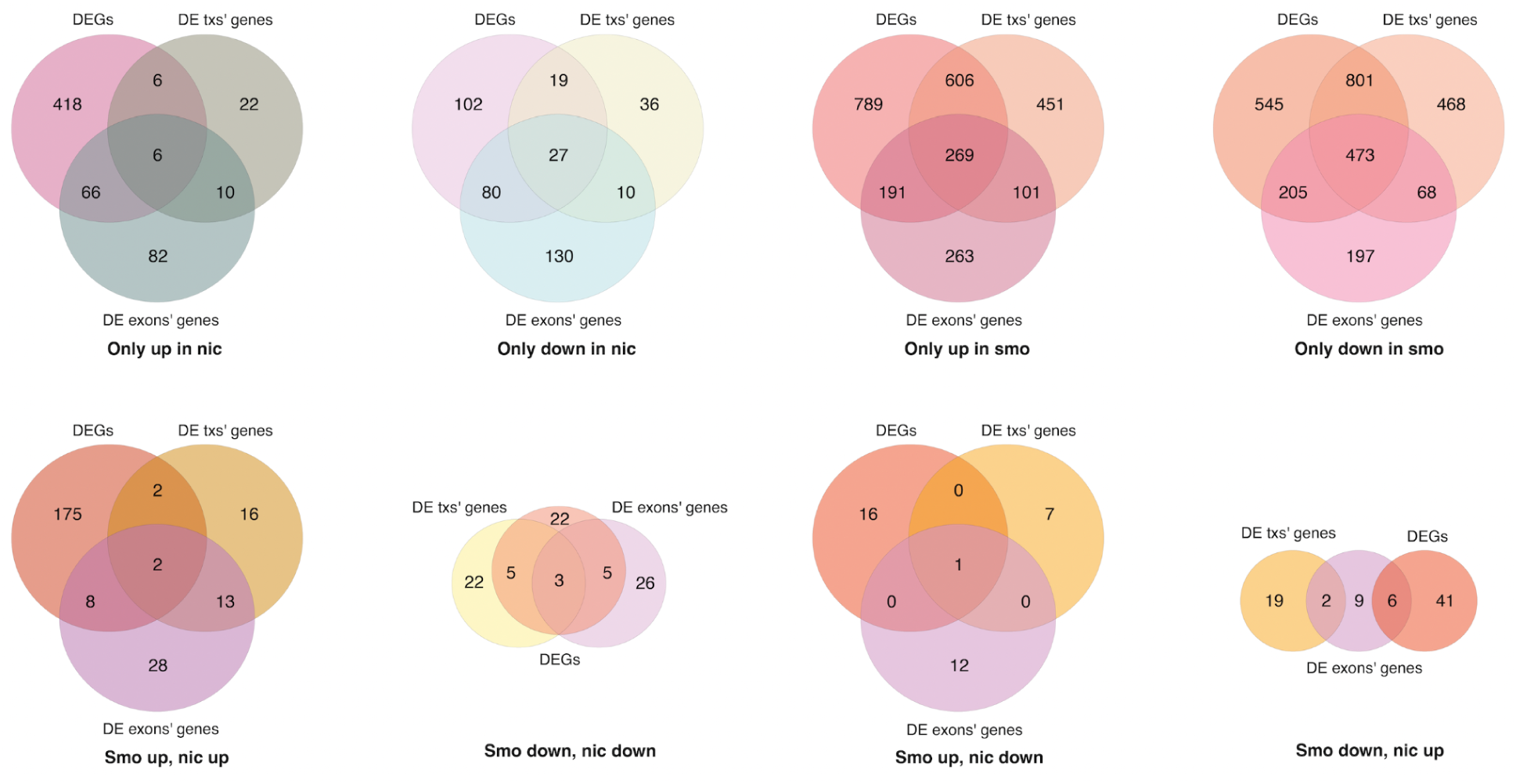
Comparison of DEA results at gene, transcript, and exon levels in pup brain. Number of DEGs and genes of DE exons and DE transcripts (txs) are shown for the groups of up- and down-regulated features for the smoking and nicotine exposures as in **Fig. S14**.

**Supplementary Figure 21:**
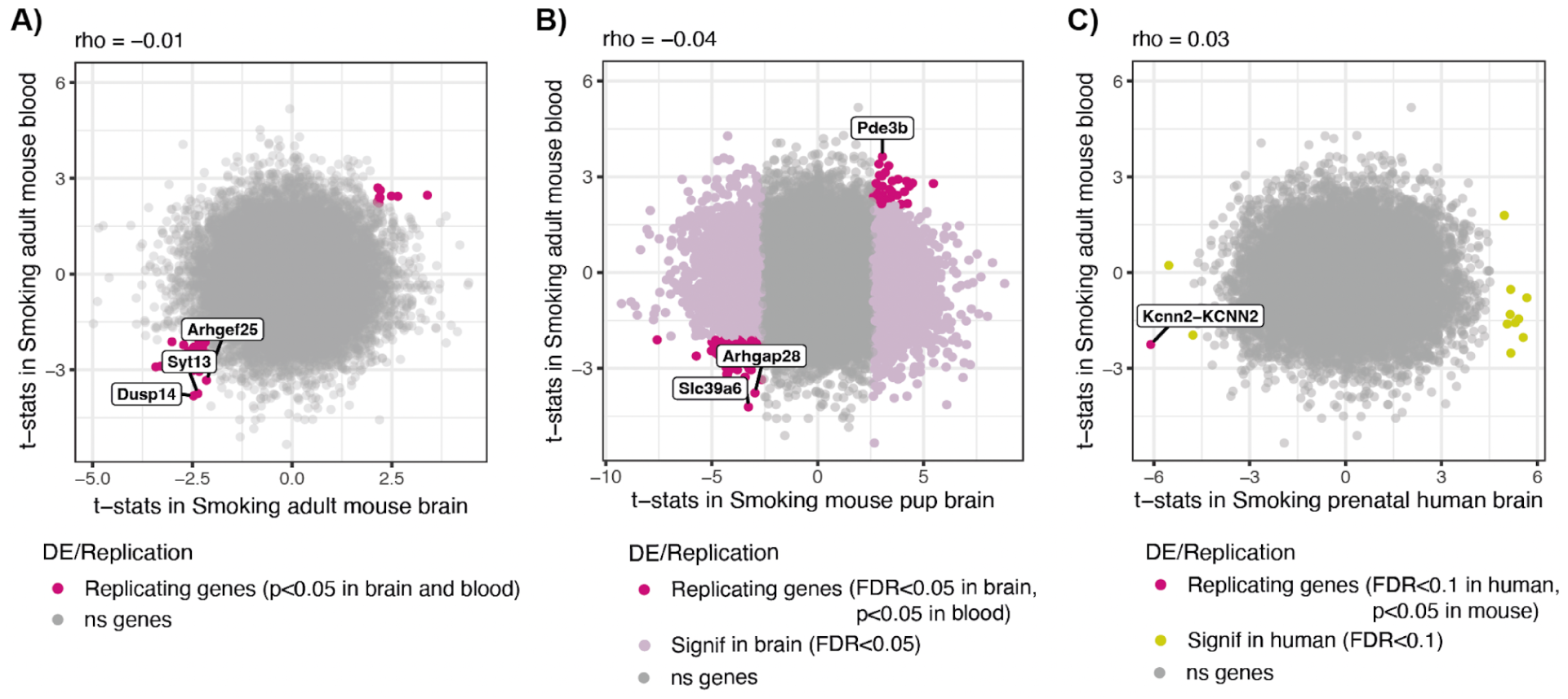
Differential gene expression signal for smoking exposure in brain and blood. Comparison of the moderated *t*-statistics of the genes for DE by smoking exposure in adult blood vs **A)** adult brain, **B)** pup brain, and **C)** prenatal human brain. In dark pink genes that replicate in blood (with *p*-value<0.05 in adult brain/ FDR<0.05 in pup brain/ FDR<0.1 in human brain, *p*-value<0.05 in blood and with same logFC sign in both tissues). For B), in light pink DEGs in pup brain (FDR<0.05). For C), in yellow DEGs in human brain (FDR<0.1). In gray genes that were non-DEGs. The three replicating genes most significant in blood are labeled with their symbol in A) and B) and the unique replicating human gene in C) is labeled together with the symbol of the orthologous gene in mouse. Rho is the Spearman correlation coefficient between the *t*-stats. Related to **Table S5** and **Table S16**.

## Supplementary File 1: additional DTE, DEE, and DJE results

### Findings beyond the gene level for gestational nicotine and smoking exposure on developing frontal cortex

#### DTE results

In addition to the highly concordant DGE and DTE results for prenatal nicotine and smoking exposure (Fig. 3A), nicotine exposure caused changes in the expression levels of transcripts from the non-DEGs *Phf3, Ankrd11*, *Trpc4*, and *Bcl11a*, all expressed in the brain and with relevant roles in brain development (109–113), as well as the pyroptosis gene *Scaf11* that has been used in the prognosis of low-grade gliomas (114) and associated with Parkinson’s disease

(115) (**Fig. S17**, **Table S8**). On the other hand, highly consistent with DGE results, exposure to smoking led to the downregulation of *Top2a* and *Ccnb2*, and the upregulation of *Mt2* at the gene and transcript levels (**Fig. S17**, **Table S9**). Importantly, the human ortholog of *Ccnb2* promotes cerebral ischemic stroke and lung cancer by interacting with *TOP2A* (116). Furthermore, similar to nicotine exposure, for smoking exposure there were non-DEGs such as *Btf3*, *Cyhr1*, *H13* and *Srsf6* expressing DE transcripts (**Fig. S17**, **Table S9**). Interestingly, *Btf3* is essential for in utero embryonic development (117), whereas the splicing factor gene *Srsf6* has target genes involved in brain organogenesis and is likely to be responsible of missplicing events that lead to Huntington’s disease (118,119); *Cyhr1* is known to be affected by chronic manganese (Mn) exposure that causes neurodegenerative changes in the frontal cortex (120), and the gene *H13* is crucial for embryonic development and brain morphology (121).

One interesting aspect was the identification of DE transcripts regulated in an opposite direction to that of their genes, as occurred with the nicotine-exposed DEG *Dgcr8* (**Fig. S17**, **Table S8**), as well as the presence DEGs with both up- and down-regulated DE transcripts within the same experiment, as was the case for *Pnsir* and *Dcun1d5* for nicotine, and *Meaf6*, *Ivns1abp*, *Morf4l2*, *Sin3b*, and *Ppp2r5c* for smoking exposure (**Fig. S17**, **Table S8**, **Table S9**). For these, transcripts going in the same direction as the gene accounted for a larger percentage of the total gene expression than transcripts with the opposite direction of regulation (**Fig. S17**), in line with past discoveries showing that genes tend to have dominant transcripts (122). Those genes could be subjected to differential transcript usage (DTU) in which not only their expression levels vary between conditions but also their splicing patterns change, resulting in different proportions of the expressed transcripts of a gene in one or the other condition. Future analyses of transcript expression proportions relative to the total expression of the genes will enable the inference of these events that can inform about substance exposure consequences at the transcriptional level that are disregarded by just analyzing DGE and DTE. Also, the actual posterior protein translation, post-translational modifications, and functional contribution of each individual DE transcript need to be explored.

#### DEE results

The functional enrichment analysis for genes with significantly regulated expression features further revealed non-DEGs that express DE transcripts and exons for smoking exposure that are implicated in neurotransmitter secretion and transport, regulation of neurotransmitter levels, and signal release from synapse (**Fig. S18A**), as well as products of DE transcripts of non-DEGs that present ionotropic glutamate and glutamate receptor binding activity and are part of the retromer complex, presynaptic membrane, and the presynaptic endocytic zone (**Fig. S18B,C**). Importantly, for smoking exposure there were DEGs with DE transcripts and exons whose protein products carry out their functions within synaptic vesicles and membranes (**Fig. S18C**). These additional neurological implications of gestational smoking exposure not identified by analyzing only DGE demonstrate how enriching it is to explore expression changes at these other expression feature levels (**Fig. S18**).

#### DJE results

DJE analysis was performed to find potentially novel splice isoforms, i.e., that are not annotated in GENCODE M25 (95,123). All DE exon-exon junctions except two for smoking exposure were novel, with at least one unannotated splice site or with an unknown combination of donor and acceptor sites. Of these, 5 and 201 DE junctions for nicotine and smoking exposure, respectively, were fully novel, with both splice sites unknown and without assigned gene. For these novel junctions their immediate following and preceding genes, as well as their nearest overlapping neighbor gene were located (see **Table S19**). The latter genes had a bigger overlap with the identified DEGs at the gene, transcript, and exon levels (**Fig. S22**), further supporting these nearest overlapping neighbor genes as bearers of potentially new isoforms, compared to the immediate upstream and downstream genes of DE junctions.

Together, DGE, DTE, DEE, and DJE results were concordant for both exposures in pup brain (Fig. 3C), with more highly expressed DEGs having DE transcripts, exons, and exon-exon junctions (**Fig. S19**), as well as DEGs with DE transcripts and DE exons, with all features regulated the same within each experiment (**Fig. S20**). Nonetheless, many DEGs only had significant DE signal at the gene level. One could hypothesize that these DEGs have low expression levels and thus, not enough reads to properly quantify their exons, transcripts, and junctions, but that was not necessarily the case: many of the DEGs with low mean expression didn’t have other DE expression features (blue points in **Fig. S19** with mean lognorm counts < 0), but not all DEGs without significant features had low expression values; they had smaller logFCs (blue points in **Fig. S19**). On the other hand, there were also DE features from non-DE genes (Fig. 3C), where the high expression of such genes enabled the detection of some of their features as DE (**Fig. S19**). A plausible explanation for the presence of DE exons from non-DEGs or lowly-expressed genes is that during exon quantification reads mapping to regions shared by overlapping exons were assigned to all, inflating their number of reads and artificially increasing their expression (see **Supplementary Materials and Methods: Expression quantification**).

Notwithstanding, a series of caveats must be considered at the exon and exon-exon junction levels. One main limitation of analyzing exons is the lack of consistency between exon and transcript expression and the very different methods used to estimate their expression levels. Because shared exons are part of more than one transcript, their expression levels have a different impact when they are considered separately, as independent genomic features, instead of parts of multiple transcripts with different expression levels. In the case of overlapping exons, the current approach multi-counts the reads mapping to them, as previously mentioned (19). Thus, exon expression levels, when measured independently, do not necessarily correlate with the expression of the transcripts containing them, which is estimated using a different method that attempts to probabilistically resolve read mapping ambiguities for shared and overlapping exons across transcripts (98). As a result, expression counts of exons may be inaccurate and could lead to misleading inferences when projected to the transcript level. Similar considerations apply to junction level expression counts. For future analyses, transcript assembly could offer a better alternative to revealing previously uncharacterized isoforms.

## Additional Supplementary Figures

**Supplementary Figure 22:**
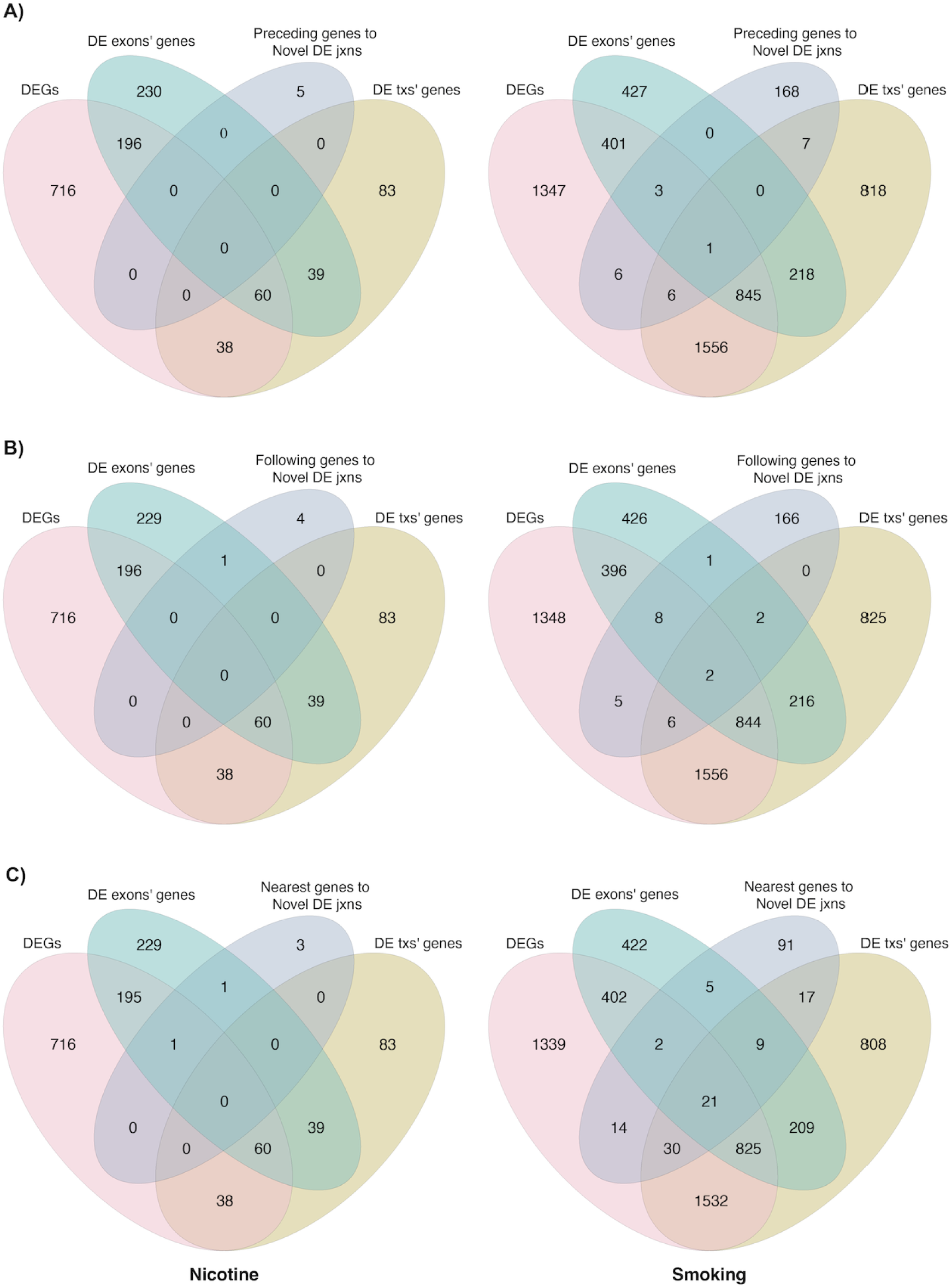
Comparison of genes DE at different feature levels and genes associated with DE exon-exon junctions in pup brain. Overlap between DEGs, genes of DE exons, genes of DE transcripts, and **A)** the preceding, **B)** following, and **C)** nearest genes to DE novel junctions without assigned gene, for nicotine (left) and smoking (right) exposure.

## Notes

https://github.com/LieberInstitute/smoking-nicotine-mouse

https://www.bioconductor.org/packages/smokingMouse/

https://www.ncbi.nlm.nih.gov/bioproject/?term=PRJNA1175674

